# Sleep and Diurnal Rest-Activity Rhythm Disturbances in a Mouse Model of Alzheimer’s Disease

**DOI:** 10.1101/2020.02.17.950600

**Authors:** Mikolaj J. Filon, Eli Wallace, Samantha Wright, Dylan J. Douglas, Lauren I. Steinberg, Carissa L. Verkuilen, Pamela R. Westmark, Rama K. Maganti, Cara J. Westmark

**Affiliations:** University of Wisconsin

**Keywords:** actigraphy, Alzheimer’s disease, CTEP, EEG, fenobam, J20 mice, mGluR_5_, sleep

## Abstract

**Study Objectives:** Accumulating evidence suggests a strong association between sleep, amyloid-beta (Aβ) deposition, and Alzheimer’s disease (AD). We sought to determine if: (1) deficits in rest-activity rhythms and sleep are significant phenotypes in J20 AD mice, (2) metabotropic glutamate receptor 5 inhibitors (mGluR_5_) could rescue deficits in rest-activity rhythms and sleep, and (3) Aβ levels are responsive to treatment with mGluR_5_ inhibitors.

**Methods:** Diurnal rest-activity levels were measured by actigraphy and sleep-wake patterns by electroencephalography (EEG), while animals were chronically treated with mGluR_5_ inhibitors. Behavioral tests were performed, and Aβ levels measured in brain lysates.

**Results:** J20 mice exhibited a 4.5 hour delay in the acrophase of activity levels compared to wild-type littermates, and spent less time in REM sleep during the second half of the light period. J20 mice also exhibited decreased NREM delta power but increased NREM sigma power. The mGluR_5_ inhibitor CTEP rescued the REM sleep deficit and improved NREM delta and sigma power but did not correct rest-activity rhythms. No statistically significant differences were observed in Aβ levels, rotarod performance or the passive avoidance task following chronic mGluR_5_ inhibitor treatment.

**Conclusions:** J20 mice have disruptions in rest-activity rhythms and reduced homeostatic sleep pressure (reduced NREM delta power). NREM delta power was increased following treatment with an mGluR_5_ inhibitor. Drug bioavailability was poor. Further work is necessary to determine if mGluR_5_ is a viable target for treating sleep phenotypes in AD.

**Statement of Significance:** Sleep disruption is evolving as an important risk factor as well as phenotype of neurological diseases including Alzheimer’s disease. This study is novel in determining alterations in the rest-activity rhythm and sleep-wake pattern of J20 Alzheimer’s disease mice and wild type littermates. Specifically, there is a delay in acrophase with prolonged hyperactivity during the dark cycle, and reduced sleep pressure that was improved by treatment with mGluR_5_ inhibitor. Critical remaining knowledge gaps and future directions include testing the effects of Alzheimer’s disease drugs on rescue of sleep and rest-activity patterns in other Alzheimer’s disease models. These studies are relevant to human Alzheimer’s disease as monitoring sleep phenotypes may predict disease risk, and therapies that normalize sleep patterns may slow progression.

## Introduction

Alzheimer’s disease (AD) is the sixth most common cause of death in the United States, afflicting approximately 5.4 million Americans, and presents a tremendous emotional and financial hardship on patients and caregivers. AD is a progressive form of dementia characterized histologically by amyloid-beta (Aβ) plaques, neurofibrillary tangles and neuronal cell death. In a small percentage of cases, AD is directly associated with specific genetic mutations in amyloid beta protein precursor (AβPP) (chromosome 21), presenilin 1 (chromosome 14) or presenilin 2 (chromosome 1); however, in the vast majority of cases the cause of the disease is unknown. Patients suffer memory loss, impaired judgment, cognitive dysfunction, the inability to perform everyday tasks, and behavioral problems. There are currently no cures for AD, which provides a strong impetus to discover novel therapeutic strategies for treatment and improved outcome measures to bridge preclinical and clinical research.

Deterioration of rest-activity cycles is a progressive phenotype in AD patients in whom reported sleep disturbances include increased nocturnal awakenings, decreased duration of REM sleep, and diminished slow-wave sleep.^1–5^ There is now evidence that rest-activity rhythm fragmentation and sleep disturbances may precede the onset of AD and drive disease pathology.^6, 7^ Restlessness, agitation, irritability and/or confusion worsen in the late afternoon and evening and last into the night with less pronounced symptoms earlier in the day. Thus, we asked if an AD mouse model exhibited altered diurnal rest-activity patterns as determined by actigraphy, assessed electroencephalogram (EEG)-based sleep-wake patterns, and determined whether any aberration in AD mice could be rescued by modulation of metabotropic glutamate receptor 5 (mGluR_5_) signaling.

Two classes of drugs, cholinesterase inhibitors (donepezil, rivastigmine and galantamine) and NMDA receptor antagonists (memantine) are currently approved by the FDA to treat cognitive symptoms of AD. These drugs act on healthy neurons to compensate for lost acetylcholine activity or modulate NMDA receptor activity, respectively. They improve cognitive ability for a year or less, but do not reduce Aβ or neurofibrillary tangle accumulation and subsequent disease progression. Aβ immunotherapy has proven to be very effective in reducing soluble Aβ, amyloid plaque and soluble tau as well as associated cognitive decline; however, there are questions about safety and it is only experimental at this point.^8–11^

An alternative, viable therapeutic target for the treatment of AD may be metabotropic glutamate receptor 5 (mGluR_5_) inhibitors. There is a strong rationale for studying mGluR_5_ inhibitors in AD models. mGluR_5_ is a glutamate-activated, G-protein-coupled receptor widely expressed in the central nervous system (CNS) and clinically investigated as a drug target for a range of indications including depression, Parkinson’s disease and fragile X syndrome (FXS). APP synthesis is regulated through an mGluR_5_-dependent signaling pathway.^12, 13^ The knockout of *mGluR_5_* in APP_SWE_/PS1ΔE9 AD mice reduces spatial learning deficits, Aβ oligomer formation and Aβ plaque number.^14^ Treatment with mGluR_5_ inhibitors reduces APP and Aβ levels and improves memory and cognitive function in mouse models.^15–17^ Herein, we test the effects of mGluR_5_ inhibition on rest-activity rhythms, sleep, locomotor ability, learning and memory, and Aβ levels in J20 mice.

The J20 mouse model is an established rodent model for the study of AD that expresses the human amyloid protein precursor (*hAPP*) gene containing both the Swedish and Indiana familial mutations. J20 mice exhibit greatly exacerbated Aβ production and cognitive deficits.^18^ The inclusion of flanking sequences in the transgenic construct is expected to affect posttranscriptional regulation of the *APP* gene and more closely mimic normal temporal and spatial expression of APP and metabolites.^19^ Herein, we show that J20 mice exhibited a pronounced 4.5 hour shift in acrophase (peak activity levels) during the dark phase of the diurnal cycle, and reduced sleep homeostatic pressure as measured by NREM delta power. Treatment with mGluR_5_ inhibitors did not change rest-activity rhythms but CTEP improved NREM delta power.

## Methods

### Mouse husbandry

The J20 [B6.Cg-Tg(PDGFB-APPSwInd)20Lms/2Mmjax] mouse model of AD expresses a mutant version of *hAPP* carrying both the Swedish (K670N/M671L) and the Indiana (V717F) mutations directed by the human PDGFB promoter. Hemizygous male J20 mice were purchased from Jackson Laboratories (catalog #006293) and mated with C57BL/6J female mice (Jackson Laboratories, catalog #000664) to generate J20 and wild type (WT) littermates. Mice were group housed in microisolator cages on a 6am-6pm light cycle with *ad libitum* access to food (Teklad 2019) and water. Mouse ages and treatments for specific experiments are defined in the figure legends. The bedding (Shepherd’s Cob + Plus, ¼ inch cob) contained nesting material as the only source of environmental enrichment. All animal husbandry and euthanasia procedures were performed in accordance with National Institutes of Health and an approved University of Wisconsin-Madison IACUC animal care protocol. J20 genotypes were determined by PCR analysis of DNA extracted from tail biopsies with HotStarTaq polymerase (Qiagen, catalog #203205) and Jackson Laboratories’ primer sequences oIMR2044 (transgene forward; 5’-GGT GAG TTT GTA AGT GAT GCC-3’) and oIMR2045 (transgene reverse; 5’-TCT TCT TCT TCC ACC TCA GC-3’) targeted at the APP_SW/IND_ transgene (360 base pair PCR product) and oIMR8744 (internal positive control forward; 5’-CAA ATG TTG CTT GTC TGG TG-3’) and oIMR8745 (internal positive control reverse; 5’-GTC AGT CGA GTG CAC AGT TT-3’), which produce an internal positive control PCR product of 200 base pair. J20 mice exhibited a premature mortality phenotype (**Figure S1**), which is consistent with prior studies in Tg2576.^20^

### Drug Preparation & Chronic Dosing

(Method 1, adult mice) The drugs fenobam (gift from FRAXA Research Foundation), 37.5 mg, and CTEP (2-chloro-4-((2,5-dimethyl-1-(4-(trifluoromethoxy)phenyl)-1H-imidazol-4-yl)ethynyl)pyridine) (MedChem Express, catalog #HY-15445), 5 mg, were transferred into an IKA Ball-Mill Tube BMT-20-S containing 10 stainless steel balls with 7.5 mL 0.9% NaCl, 0.3% Tween-80. Drug plus vehicle were mixed on low velocity for 1 min and high velocity setting #9 for 5 min. Additional vehicle (7.5 mL) was added and the drug was mixed on high velocity setting #9 for 5 min. Fenobam stock was 2.5 mg/mL. CTEP was further diluted with 9.75 mL vehicle resulting in final drug concentrations of 0.2 mg/mL CTEP. Vehicle and drugs were frozen in single-use aliquots at −20^0^C. Mice were dosed once daily with fenobam and once every other day with CTEP at 400 μL per 40g body weight by oral gavage with 22g 1.4” feeding needles with ball (Kent Scientific, catalog #FNC-22-1.5). Final drug concentrations were 24 mg/kg fenobam and 2 mg/kg CTEP. Mice were typically dosed midway through the light cycle (11am-1pm). (*Method 2, aged mice*) CTEP (10 mg) was dissolved in 200 μL DMSO and aliquots at 50 mg/mL were frozen at −20^0^C. On the day of use, 40 μL of CTEP was mixed with vehicle [1 wt% Hypromellose (HPMC) (Sigma catalog #H3785) and 1 wt% Tween-80] in an IKA Ball-Mill Tube BMT-20-S containing 10 stainless steel balls as described above to a final concentration of 0.2 mg/mL CTEP. Mice were dosed at 2 mg/kg by oral gavage.

### Neuroassessments

Mice underwent an abbreviated Irwin murine neurobehavioral screen, including weekly weight measurements, at the beginning and end of the drug dosing regimen (**Tables S1 & S2**).^21–23^

**Table 1:** Mouse Cohorts for Actigraphy Experiments. Three cohorts of wild type (WT) and J20 littermate mice were tested by actigraphy during various seasons (set A: fall, set B: winter, set C: spring). A minimum of 8 mice were tested per cohort. Average age of the mice was 8 months old (presented in days ± the standard deviation). Average peak acrophase is in minutes ± standard deviation. <u>2-way ANOVA based on season and genotype</u>: interaction *p*=0.033, F(2,81)=3.56; season *p*=0.57, F(2,81)=0.57; genotype, *p*<0.0001, F(1,81)=49.8.

| **Table 1: Mouse Cohorts for Actigraphy Experiments** |  |  |  |  |  |  |
| --- | --- | --- | --- | --- | --- | --- |
| **Experiment** | **WT** |  |  | **J20** |  |  |
|  | **N** | **Age (days)** | **Peak Acrophase (min)** | **N** | **Age (days)** | **Peak Acrophase (min)** |
| A: fall | 15 | 231±8 | 836±82 | 13 | 232±8 | 1081±143 |
| B: winter | 20 | 248±13 | 941±142 | 19 | 244±13 | 1041±121 |
| C: spring | 12 | 260±5 | 859±117 | 8 | 259±7 | 1096±104 |

### Actigraphy

Rest-activity rhythms were assessed under standard lighting conditions in home-made Plexiglas® chambers containing passive infrared sensors mounted on the underside of the lids.^24, 25^ The dimensions of the transparent cylindrical Plexiglas® chambers were 6” diameter X 10” height. Mice were individually housed during actigraphy with access to food and water. Each gross movement of the animal was recorded as an activity count with VitalView acquisition software (Minimitter Inc., Bend, OR, USA). Activity counts were binned in 60 second epochs and scored on an activity scale (0-50) over a 3-9 day period. Data were analyzed with ACTIVIEW Biological Rhythm Analysis software (Mini Mitter Company, Inc.). A Chi-square periodogram method was used to determine the diurnal rest-activity period.

### EEG Sleep Analysis

Mice (age 11-12 months old) were recorded in sleep-wake patterns using electroencephalographic (EEG) monitoring. For EEG electrode implantation surgery (Day 1), anesthesia was induced with 5% isoflurane and maintained at 1-2% in oxygen flowing at 0.5-1 L per minute. Three stainless steel epidural screws were placed as electrodes with two screws over the frontal (Bregma +1.5 mm and +1 mm laterally) and parietal cortex (Bregma −3 mm and − 1 mm laterally) and one occipital reference (lambda −1 mm at midline). Two stainless steel wire electrodes were placed in the nuchal muscles for electromyography (EMG) recording. The EEG and EMG electrodes were connected to a head cap and secured with dental acrylic. Standard analgesia was administered per local IACUC recommendations. Mice were allowed to recover from the surgery (Days 2 & 3, singly housed) prior to transfer to individual, tethered EEG acquisition chambers and dosing with CTEP (Days 4, 6, 8 and 10). EEG recordings and analyses have been previously described.^25, 26^ Recordings were acquired Days 8-12 on an XLTEK machine (Natus, Madison, WI) with a 512 Hz sampling rate, and the three full days of recordings (Days 9-11) were used for the analysis. EEG recordings were manually scored in 4-second epochs for REM, NREM and awake vigilance states with Sirenia Sleep software v.2.0.4 by scorers blinded with respect to treatment group. Waking epochs were identified as those with high EMG amplitude (**Figure S2A**). Epochs with relatively quiescent EMG were scored as sleep. Specific sleep states were differentiated based on predominant EEG power such that NREM was associated with high amplitude delta (1-4 Hz, **Figure S2B**) and REM was associated with low amplitude theta (5-7 Hz, **Figure S2C**) activity.

### Rotarod

The mice were acclimated to the test room for at least 20 min prior to testing on a Rotarod Treadmill (Med Associates Inc., Vermont, USA). The rotarod was set to a speed setting of 9, which accelerates from 4.0-40 rpm over 5 min. Mice were placed on the rotarod and the latency time to when the mouse fell off was recorded. If a mouse made two complete turns hanging onto the grip bar without actively walking/running, the mouse was counted as falling off of the beam. If more than 300 seconds elapsed, the mouse was removed from the beam. Experiments entailed 4 trials on day 1 and 2 trials on day 2.

### Passive Avoidance

Mice were acclimated to the experimental room for at least 20 minutes prior to testing in a foot shock passive avoidance paradigm using an aversive stimulator/scrambler (Med Associates Inc., Vermont, USA). A bench-top lamp was turned on behind the center of a light/dark shuttle box and aimed toward the back-left corner away from the dark side of the shuttle box. The power supply on the shock grid was set at 0.6 mA. The apparatus was cleaned with 70% ethanol between animals. On the training day, a mouse was placed in the light side of the shuttle box toward the back corner away from the opening to the dark side of the shuttle box. The trap door in the shuttle box was open. After the mouse crossed over to the dark side, the trap door was closed and the latency time for the mouse to move from the light to the dark side was recorded. The mouse was allowed to equilibrate in the dark side for 5 seconds before receiving a 2-second 0.6 mA footshock. After 15 seconds, the mouse was removed from the shuttle box and returned to its home cage. At test times, the mouse was placed in the light side of the shuttle box facing the left rear corner away from the opening to the dark side with the trap door open. The latency time for the mouse to move from the light to the dark side was recorded. If the mouse did not move to the dark side within 300 seconds, it was gently guided to the dark side. The mouse was allowed to equilibrate to the dark side for 5 seconds before return to the home cage. Mice were tested 6, 24, and 48 hours after training. Mice only received one shock on the training day. Testing at 24 and 48 hours measured extinction.

### Tissue Collection

Mice were treated with isoflurane for 1 minute and blood collected from the abdominal artery with a 21Gx¾”x12” vacutainer blood collection set (Becton Dickinson, catalog #367296). The blood was immediately mixed with sodium heparin (20 μL of 10 mg/mL; Sigma #H3393). Brain tissue (hippocampus, cerebellum and right and left cortices) were dissected and quick frozen on dry ice. Tissue was collected to confirm genotype analyses. The heparinized blood was spun for 10 minutes at 5,000 rpm at room temperature and the upper plasma layer was transferred to a 1.5 mL Eppendorf tube, frozen on dry ice and stored at −80^0^C. *Pharmacokinetics*: Plasma and brain left cortices were shipped to Tandem Labs (Durham, NC) for detection of CTEP levels by mass spectrometry analysis. Study samples were analyzed using standards prepared in sodium heparin mouse plasma. The method calibration range was 0.500-10,000 ng/mL using two different transitions for CTEP. The C13 peak for CTEP was used for the top 5 calibration points and the C12 peak for CTEP was used for the bottom 5 points of the curve. CTEP and fenobam stock solutions for the calibration curve and internal standard, respectively, were prepared at 1 mg/mL in 50:50 water:acetonitrile.

### Brain Lysates & Aβ ELISA

Diethylamine (DEA) protein extraction buffer [20 μL DEA (0.2% final; Fisher catalog #A11716), 0.5 mL 1M NaCl (50 mM final), 2 mL 10X protease inhibitor cocktail (RPI catalog #P50600) in a 10 mL final volume] was chilled in ice. Tissue to be homogenized (right cortex of brain) was transferred to a Dounce glass-glass homogenizer with 5 volumes ice-cold DEA protein extraction buffer (1 mL per 200 mg tissue) and homogenized with 35 strokes.

Lysates were spun at 20,000g for 30 min at 4^0^C. The cleared supernatant was removed and neutralized with 1/10 volume 0.5M Tris, pH 6.8. Aliquots were quick frozen at −80^0^C and protein concentrations quantitated by the BCA Assay (Pierce, catalog #23235) per the manufacturer instructions. Aβ_1-40_ and Aβ_1-42_ levels were quantitated with Wako Human/Rat Aβ40 (catalog #294-64701) and Wako Human/Rat Aβ_1-42_ high sensitivity (catalog #292-64501) ELISA kits per the manufacturer instructions. Plasma samples were diluted 4-fold with standard diluent buffer containing protease inhibitor cocktail to disrupt interactions between Aβ with masking proteins. Brain samples were diluted 1:50 (WT) and 1:500 (J20) with standard diluent. Antibody-coated plates were incubated with standards and samples overnight at 4^0^C.

### Statistical Methods

Statistical significance for the actigraphy comparing two groups was determined by two-sided T-tests with Bonferroni corrections (1,440 time points, alpha value of 0.0000347) using Excel v16.21 software. Black dots across the top of the actigraphy graphs represent statistical significance. Statistical significance for peak acrophase and ELISAs was determined by two-way ANOVA using GraphPad Prism v8.3.0 for Mac OS X (GraphPad Software, San Diego, CA) to compare the means of 3 or more unmatched groups. Statistical analyses for EEG-based experiments utilized Matlab software (The MathWorks, Inc., Natick, MA). EEG power analysis compared normalized power of delta, theta, sigma, and gamma frequency bands of NREM sleep bouts as determined by manual scoring. As with sleep scoring, all frequency bands were calculated from 4-second epochs of 60 Hz notch-filtered EEG, and grouped within 2-hour segments across the light cycle. Sleep efficiency was assessed as percent-time in each vigilance state (i.e., wake, NREM, and REM) in four 6-hour bins. Both were evaluated using a mixed-model N-way ANOVA using group (i.e. WT-Vehicle, J20-CTEP) as a fixed-effect variable, and time as a random-effect variable. Each group included in ANOVA analysis was tested for skewness and satisfied normality with values less than |2|. Statistical analyses were conducted with an alpha value of 0.05. Cohort sizes are listed in the figure legends and means and SEM or 95% confidence intervals (CI) are graphed.

## Results

### Rest-Activity Rhythms are Disrupted in J20 Mice

Three independent experiments were performed on cohorts of WT and J20 littermate mice at 8 months of age to assess the effect of genotype and season on rest-activity patterns (**Figure 1, Table 1**). There was a highly reproducible delay in peak acrophase during the dark cycle in the J20 mice irrespective of season. Specifically, WT mice exhibited peak activity between 7-8 pm and J20 exhibited peak activity between 11pm-12am resulting in an approximately 4.5 hour delay in acrophase in J20 mice. The findings were consistent in three independent sets of data that compared testing during the fall, winter and spring, albeit the differences were the most pronounced during the spring followed by the fall and winter. On periodograms, the diurnal period was similar between WT (24.3±0.8; n=8) and J20 (24.5±0.9; n=11) mice (*p*=0.63). Genotype did not have a significant effect on daytime activity although J20 mice exhibited increased activity (11%) during the first hour in the actigraphy chambers, which likely represents decreased habituation to a novel environment (**Table 2; Figure S3**).

**Figure 1:**
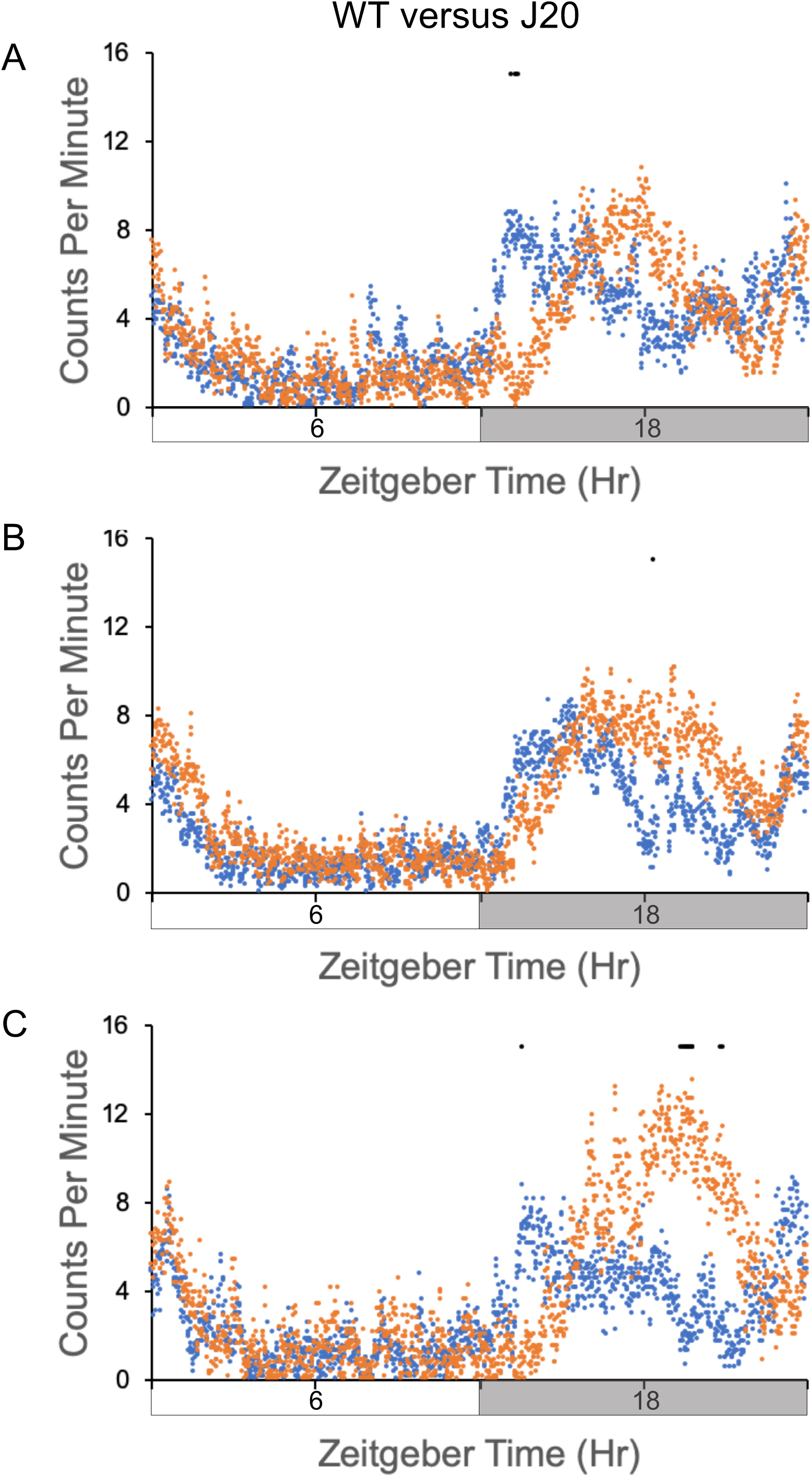
J20 mice exhibit delayed acrophase during the dark cycle. Activity counts on day 1 in the actigraphy chambers were assessed in 3 separate cohorts of wild type (WT) (blue) and J20 (orange) 8-month old mice (A=fall, B=winter, C=spring). Total activity counts (binned in 1 minute increments) were averaged for cohorts and plotted on the y-axis versus a 24 hour time period (in minutes) on the x-axis. Time zero is “Lights On”. (A) Cohort 1 consists of WT (n=15) and J20 (n=13). (B) Cohort 2 consists of WT (n=20) and J20 (n=19). (C) Cohort 3 consists of (n=12) and J20 (n=8). Black dots at the top of the graphs represent statistical significance determined by T-test with Bonferroni correction.

**Table 2:** Activity Counts During Habituation to the Actigraphy Chambers. Activity counts during the first 60 min in the novel environment of the actigraphy chambers were assessed in wild type (WT) and J20 cohorts for the 3 independent experiments. A minimum of 8 mice were tested per cohort. The average age of the mice was 8 months old. Average activity counts are presented ± the standard deviation.

| **Table 2: Activity Counts During Habituation to the Actigraphy Chambers** |  |  |  |  |  |
| --- | --- | --- | --- | --- | --- |
| **Experiment** | **WT** |  | **J20** |  | **TTEST** |
|  | **N** | **Activity Counts** | **N** | **Activity Counts** | **P value** |
| A: fall | 15 | 737±122 | 13 | 818±73 | <0.05 |
| B: winter | 12 | 763±64 | 11 | 832±61 | <0.02 |
| C: spring | 12 | 767±71 | 8 | 860±38 | <0.004 |

### mGluR_5_ Inhibitors Do Not Restore Rest-Activity Rhythms in J20 Mice

We then tested whether the mGluR_5_ inhibitors fenobam and CTEP could restore typical rest-activity rhythms. We also performed a behavioral battery along with the actigraphy (**Figure S4**). Mice underwent a pre-treatment evaluation of general fitness and grip strength as previously described^21^ as well as weekly assessments throughout dosing (**Table S1** (fenobam) and **Table S2** (CTEP)). The mGluR_5_ inhibitors were administered by oral gavage either daily (fenobam) or every other day (CTEP). Neither WT nor J20 mice exhibited alterations in general fitness resulting from treatment. No differences were seen in rest-activity patterns with fenobam (**Figure 2**). Peak acrophase in the WT and J20 cohorts [WT/vehicle 912±84 min, WT/fenobam 940±84 min, J20/vehicle 1,050±87 min, J20/fenobam 1,007±97min; 2-way ANOVA: interaction *p*=0.30, F(1,24)=1.11; drug treatment *p*=0.81, F(1,24)=0.06; genotype, *p*<0.005, F(1,24)=9.52] was similar to the data for the untreated mice in **Table 1**. Likewise, CTEP did not alter peak acrophase in WT mice; however, the stress of chronic injections muted the difference between WT and J20 mice, which was restored with CTEP (**Figure 3**) [WT/vehicle 1,002±96 min, WT/CTEP 1,020±104 min, J20/vehicle 1,108±137 min, J20/CTEP 1,094±94 min; 2-way ANOVA: interaction *p*=0.61, F(1,43)=0.26; drug treatment *p*=0.95, F(1,43)=0.0048; genotype, *p*<0.0062, F(1,43)=8.29].

**Figure 2:**
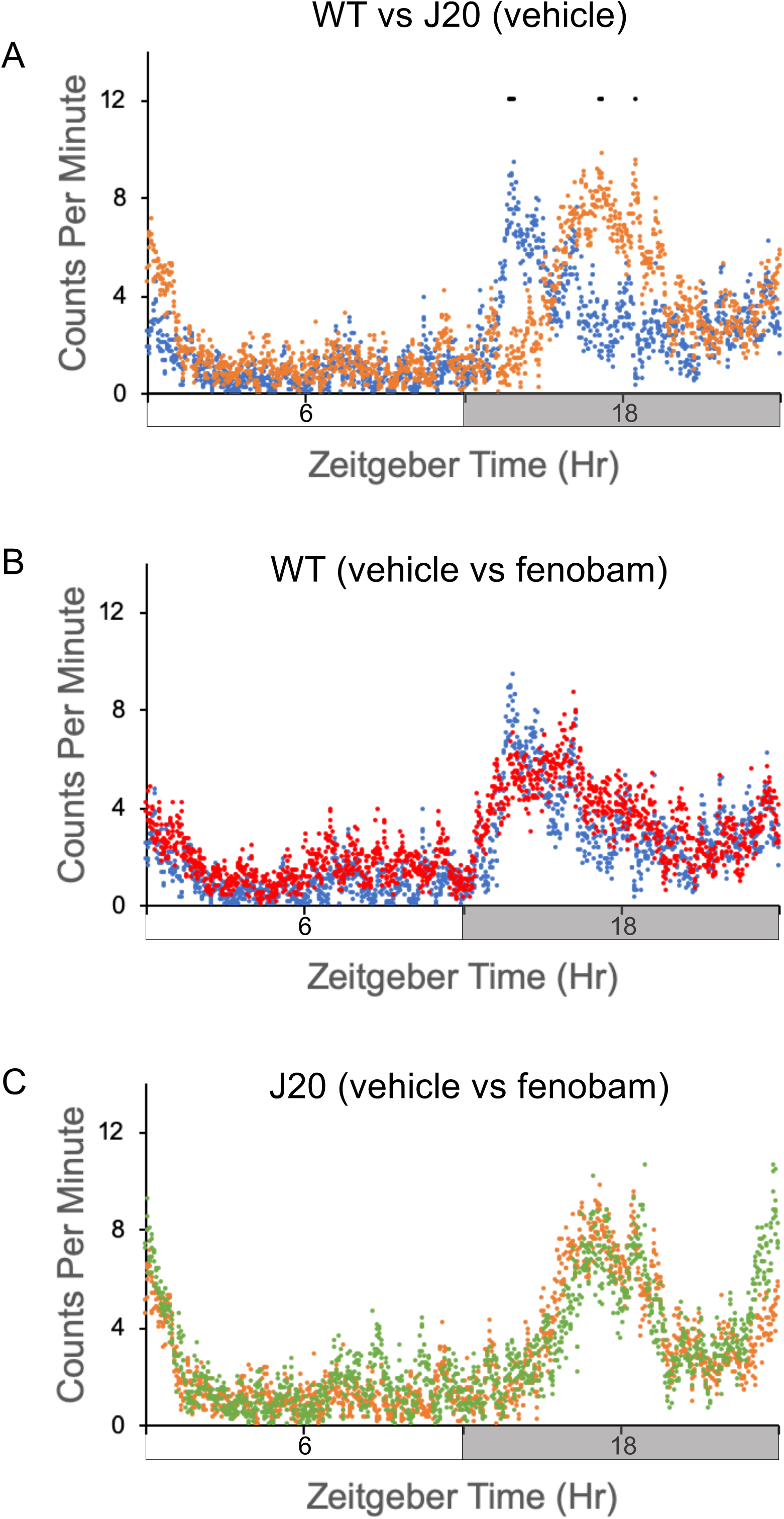
Diurnal activity levels in wild type (WT) and J20 mice in response to fenobam. Activity counts were assessed in 8-month old WT and J20 mice after chronic treatment with vehicle or fenobam. Total activity counts (binned in 1 minute increments) were averaged over 3-4 days of readings for cohorts and plotted on the y-axis versus a 24 hour time period (in minutes). Time zero is “Lights On”. Cohorts consist of WT mice treated with vehicle (n=7), J20 treated with vehicle (n=7), WT treated with fenobam (n=8), and J20 treated with fenobam (n=6). (A) vehicle-treated WT (blue) versus J20 (orange). (B) WT mice treated with vehicle (blue) versus fenobam (red). (C) J20 mice treated with vehicle (orange) versus fenobam (green). Black dots at the top of the graphs represent statistical significance determined by T-test with Bonferroni correction. 2-way ANOVA results: interaction *p*<0.0001, F(4317, 34560)=1.73; treatment (genotype and drug) *p*<0.0001, F(3, 34560)=155.8; time, *p*<0.0001, F(1439, 34560)=10.25.

**Figure 3:**
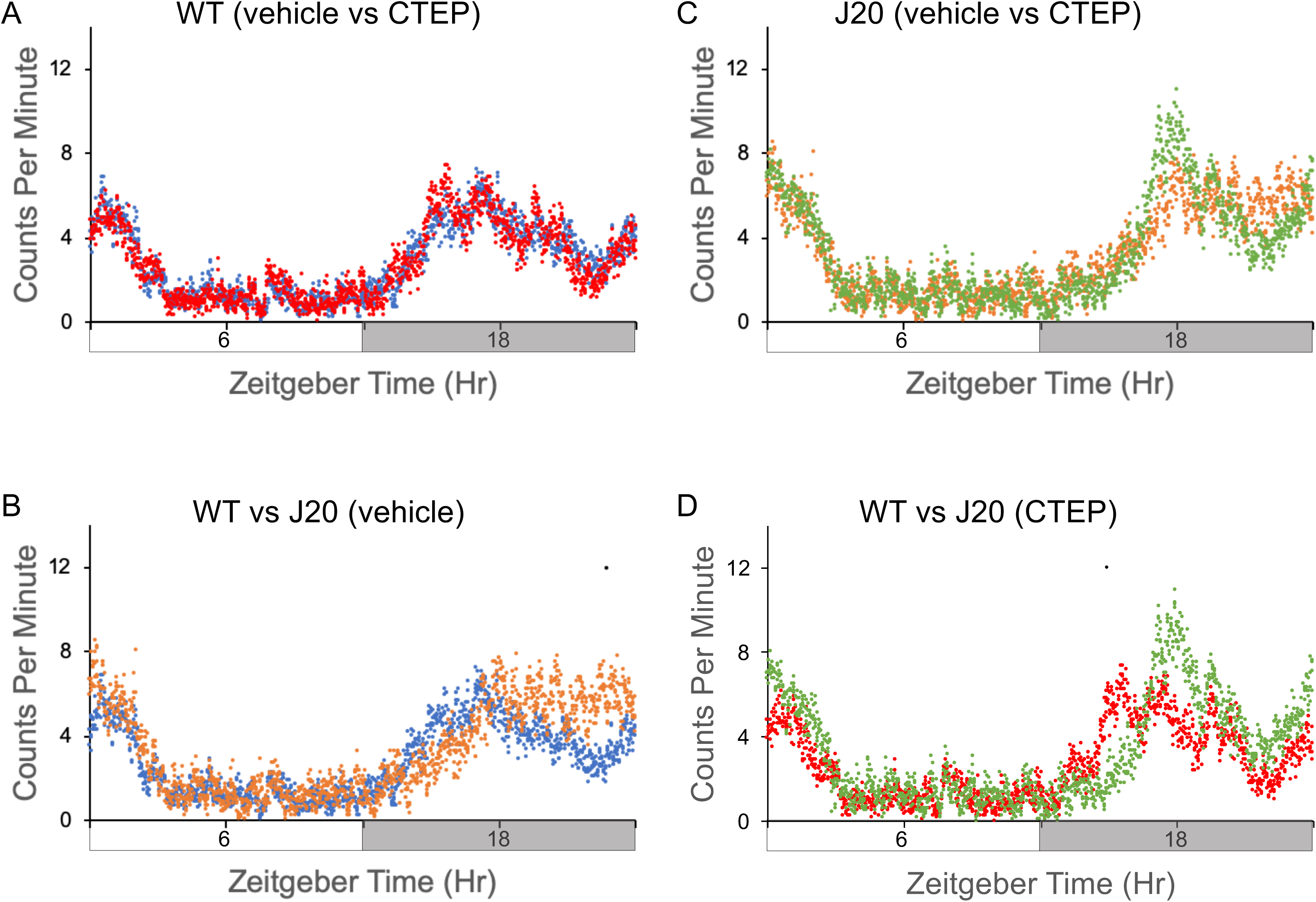
Diurnal activity levels in wild type (WT) and J20 mice in response to CTEP (2-chloro-4-((2,5-dimethyl-1-(4-(trifluoromethoxy)phenyl)-1H-imidazol-4-yl)ethynyl)pyridine). Activity counts were assessed in 9-month old WT and J20 mice after chronic treatment with vehicle or CTEP. Total activity counts (binned in 1 minute increments) were averaged over 2-4 days of readings for cohorts and plotted on the y-axis versus a 24 hour time period (in minutes). Time zero is “Lights On”. Cohorts consist of WT mice treated with vehicle (n=14, blue), J20 treated with vehicle (n=9, orange), WT treated with CTEP (n=12, red), and J20 treated with CTEP (n=12, green). (A) WT mice treated with vehicle (blue) versus CTEP (red). (B) J20 mice treated with vehicle (orange) versus CTEP (green). (C) vehicle-treated WT (blue) versus J20 (orange). (D) CTEP-treated WT (red) versus J20 (green). Black dots at the top of the graphs represent statistical significance determined by T-test with Bonferroni correction. 2-way ANOVA results: interaction *p*<0.0001, F(4317, 61920)=1.45; treatment (genotype and drug) *p*<0.0001, F(3, 61920)=168.4; time, *p*<0.0001, F(1439, 61920)=19.19.

We also tested CTEP in aged WT and J20 mice (16-19-month old mice). Neither genotype nor CTEP statistically altered peak acrophase in aged J20 mice (2-way ANOVA: interaction *p*=0.62, F(1,25)=0.25; drug treatment *p*=0.43, F(1,25)=0.64; replicates, *p*=0.056, F(1,25)=4.03). (**Figure S5**). Average total daily activity counts were not statistically different between WT and J20 mice irrespective of treatment, albeit there were trends for increased activity counts in the J20 mice (**Table 3**). It should be noted that studies in the J20 mice represent the survivors as there was a premature mortality phenotype in the animals (**Figure S1**). Overall, chronic dosing with fenobam and CTEP did not rescue altered rest-activity rhythms in J20 mice.

**Table 3:** Activity Levels in Response to mGluR5 Treatment. Total daily average activity counts ± standard deviation were assessed in wild type (WT) and J20 mice in response to vehicle, fenobam and CTEP (2-chloro-4-((2,5-dimethyl-1-(4-(trifluoromethoxy)phenyl)-1H-imidazol-4-yl)ethynyl)pyridine) in 8 month old mice and in response to CTEP in 16-19 month old mice. Fenobam was dosed daily and CTEP was dosed every other day (EOD).

| **Table 3: Activity Levels in Response to mGluR5 Treatment** |  |  |  |  |  |  |  |
| --- | --- | --- | --- | --- | --- | --- | --- |
|  | **WT** |  |  | **J20** |  |  | ***P*** |
| **Experiment** | **N** | **Age (days)** | **Activity Counts** | **N** | **Age (days)** | **Activity Counts** | **Activity Counts** |
| Vehicle (daily) | 7 | 256±11 | 3039±911 | 7 | 254±3 | 3938±880 | 0.08 |
| Fenobam (daily) | 8 | 252±3 | 3821±723 | 6 | 258±12 | 4156±715 | 0.41 |
| Vehicle (EOD) | 14 | 278±9 | 4010±784 | 9 | 276±11 | 4306±1958 | 0.62 |
| CTEP (EOD) | 12 | 277±8 | 3939±924 | 12 | 275±8 | 4347±1752 | 0.48 |
| Vehicle (EOD) | 7 | 508±38 | 3769±1082 | 7 | 504±39 | 4209±771 | 0.40 |
| CTEP (EOD) | 9 | 507±42 | 3626±1080 | 6 | 545±58 | 4304±898 | 0.23 |

### Analysis of Sleep-Wake Patterns

To qualify the effects of CTEP administration on sleep patterns, WT and J20 mice (11-12 months old) were implanted with EEG/EMG electrodes and subsequently treated with either CTEP or vehicle (n=3-4 mice/treatment cohort; treated every other day over 9 days) (**Figure 4, S6**). Acquired EEG were manually scored for vigilance states in 4-second epochs.^26^ First, 6-hour binned mixed-effect ANOVA analyses showed a main effect of time in percent time spent for each vigilance state (wake: *p*<0.01, F(3,152)=79.27 [bin mean±95CI: T1-33.49±1.79%, T2=29.35±1.79%, T3=48.83±1.79%, T4=58.14±1.79%], NREM:, *p*<0.001, F(3,152)=134.47 [bin mean±95CI: T1=58.2±1.55%, T2=60.25±1.54%, T3=44.41±1.55%, T4=35.54±1.55%], and REM: *p*<0.001, F(3,152)=51.18 [bin mean±95CI: T1=6.79±0.3%, T2=7.88±0.3%, T3=3.86±0.3%, T4=2.37±0.3%]), suggesting significant oscillation across the light-dark cycle. We also found a main effect of treatment/genotype group in percent awake time (*p*=0.03, F(3,152)=4.33 [group means±95CI: WT/vehicle=40.18±1.92%, WT/CTEP=46.64±1.66%, J20/vehicle=40.51±1.66%, J20/CTEP=42.47±1.91%]) and NREM sleep (*p*<0.01, F(3,152)=99.63 [group means±95CI: WT/vehicle=53.11±1.65%, WT/CTEP=45.88±1.4%, J20vehicle=48.0±1.43%, J20/CTEP=51.08±1.66%]). Multiple comparisons of all interactions (time*group) revealed a significant reduction in estimated marginal mean of time spent in REM sleep by vehicle-treated J20 (5.37±0.56%) relative to vehicle-treated WT (9.04±0.65%) mice during the second half of the light period, a difference not seen when J20 were treated with mGluR_5_ inhibitor (**Figure 4**).

**Figure 4:**
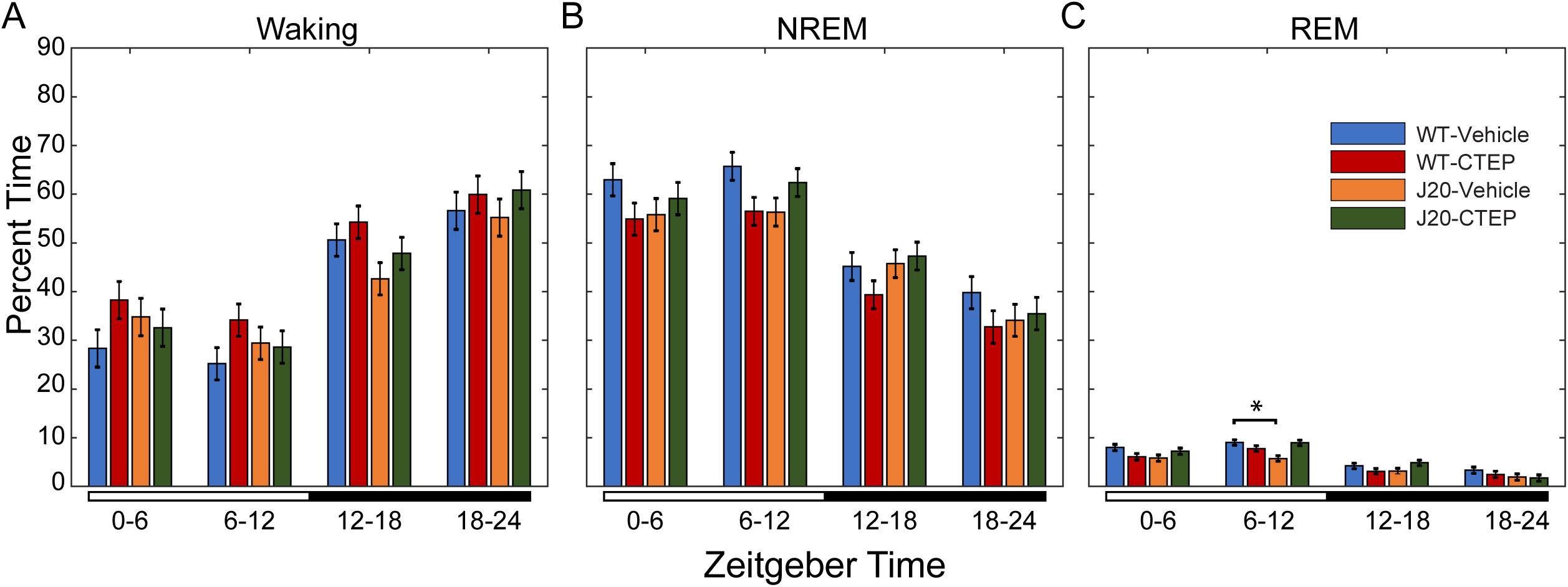
Manually scored sleep based on EEG recordings from wild-type and J20 mice with and without CTEP. Data were separated into four, 6-hour bins, starting at Zeitgeber time 0, lights-on. Lighting condition is annotated by the bar below the graphs: open: lights on and closed: lights off. Percent time in waking, and NREM and REM sleep are presented as marginal means ± 95% confidence interval (95CI) for each treatment group: wild type (WT) treated with vehicle (n=3 mice for 3 days); J20 treated with vehicle (n=4 mice for 3 days); WT treated with CTEP (2-chloro-4-((2,5-dimethyl-1-(4-(trifluoromethoxy)phenyl)-1H-imidazol-4-yl)ethynyl)pyridine) (n=4 mice for 3 days), and J20 treated with CTEP (n=3 mice for 3 days). Non-overlapping 95CI bars indicate a significant difference (*P*<0.05).

We then analyzed NREM EEG power among the different treatment groups. Each 4-second segment of 60 Hz notch-filtered EEG was evaluated with a Fast-Fourier transform, from which power of delta (0.5-4 Hz), theta (5-9 Hz), sigma (10-14 Hz), and gamma (25-100 Hz) frequencies were calculated. To account for inter-animal variability, EEG power values were normalized to the summed power of delta, theta, sigma, and gamma bands. Records were divided into 2-hour bins and EEG power of NREM epochs within each bin were grouped. Mixed model ANOVA of EEG delta power showed a main effect of time for delta (*p*<0.001, F(11, 446462)=3.92, [bin means±95CI: T1=0.63±0.00066, T2=0.612±0.00064, T3=0.599±0.00063, T4=0.578±0.00062, T5=0.569±0.00063, T6=0.575±0.00065, T7=0.564±0.00068, T8=0.579±0.00072, T9=0.587±0.00083, T10=0.589±0.00078, T11=0.602±0.00077, T12=0.625±0.00099]), theta (*p*<0.05, F(11, 446462)=1.47 [bin means±95CI: T1=0.193±0.00039, T2=0.199±0.00038, T3=0.204±0.00037, T4=0.209±0.00037, T5=0.207±0.00038, T6=0.21±0.00039, T7=0.216±0.00041, T8=0.21±0.00044, T9=0.211±0.0005, T10=0.213±0.00048, T11=0.206±0.00047, T12=0.201±0.0006]), and sigma (*p*<0.001, F(11, 446462)=4.323 [bin means±95CI: T1=0.106±0.00032, T2=0.112±0.00031, T3=0.119±0.0003, T4=0.126±0.0003, T5=0.126±0.00031, T6=0.128±0.00032, T7=0.129±0.00034, T8=0.122±0.00035, T9=0.118±0.00041, T10=0.118±0.00039, T11=0.111±0.00038, T12=0.101±0.00049]). Mixed-model ANOVA also revealed a main effect of group for delta (*p*<0.001, F(3,446462)=16.55 [group means±95ci: WT/vehicle=0.624±0.00043, WT/CTEP=0.605±0.0004, J20/vehicle=0.554±0.00039, J20/CTEP=0.587±0.00045]), theta (*p*<0.001, F(3,446462)=28.94 [group means±95ci: WT/vehicle=0.186±0.00026, WT/CTEP=0.2079±0.00024, J20/vehicle=0.219±0.00024, J20/CTEP=0.214±0.00027]), sigma (*p*<0.001, F(3,446462)=32.83 [group means±95CI: WT/vehicle=0.096±0.00021, WT/CTEP=0.124±0.0002, J20/vehicle=0.136±0.00019, J20/CTEP=0.117±0.00022]), and gamma (*p*<0.001, F(3,446462)=14.048 [group means±95CI: WT/vehicle=0.094±0.00021, WT/CTEP=0.638±0.0002, J20/vehicle=0.09±0.00019, J20/CTEP=0.083±0.00022]) frequencies.

Interaction level analysis revealed significant effects of time*group for delta (*p*<0.001, F(33,446462)=178.31), theta (*p*<0.001, F(33,446462)=92.58), sigma (*p*<0.001, F(33,446462)=103.42), and gamma (*p*<0.001, F(33,446462)=287.44) frequencies, see **Table S3** and **Figure 5** for marginal means resulting from the time*group analysis. As seen in **Figure 5A**, the oscillation magnitude of EEG delta power in vehicle treated J20 mice was significantly reduced compared to WT animals treated with vehicle, and there was a delayed dark phase rise, which was similar to the delay in acrophase determined by actigraphy (**Figure 5, Table S3**). Of note, treatment with CTEP in J20 mice resulted in a significant increase in delta power, though not to that of WT animals. CTEP appeared to have the opposite effect in WT mice where the overall and oscillation of delta power were reduced compared to vehicle-treated WT animals. Conversely, vehicle treated J20 mice exhibited consistently increased NREM EEG power in theta and sigma frequency bands (**Figure 5B and C**), which was moderately reduced in CTEP treated J20 animals and increased in CTEP treated wild-type animals. Gamma power of NREM sleep showed less consistent differences between vehicle treated WT and J20 animals. As seen in **Figure 5D**, WT animals showed significantly greater oscillation of NREM gamma power across the day. Treatment with CTEP reduced overall NREM gamma power for both genotypes, though the effect was more pronounced in WT animals.

**Figure 5:**
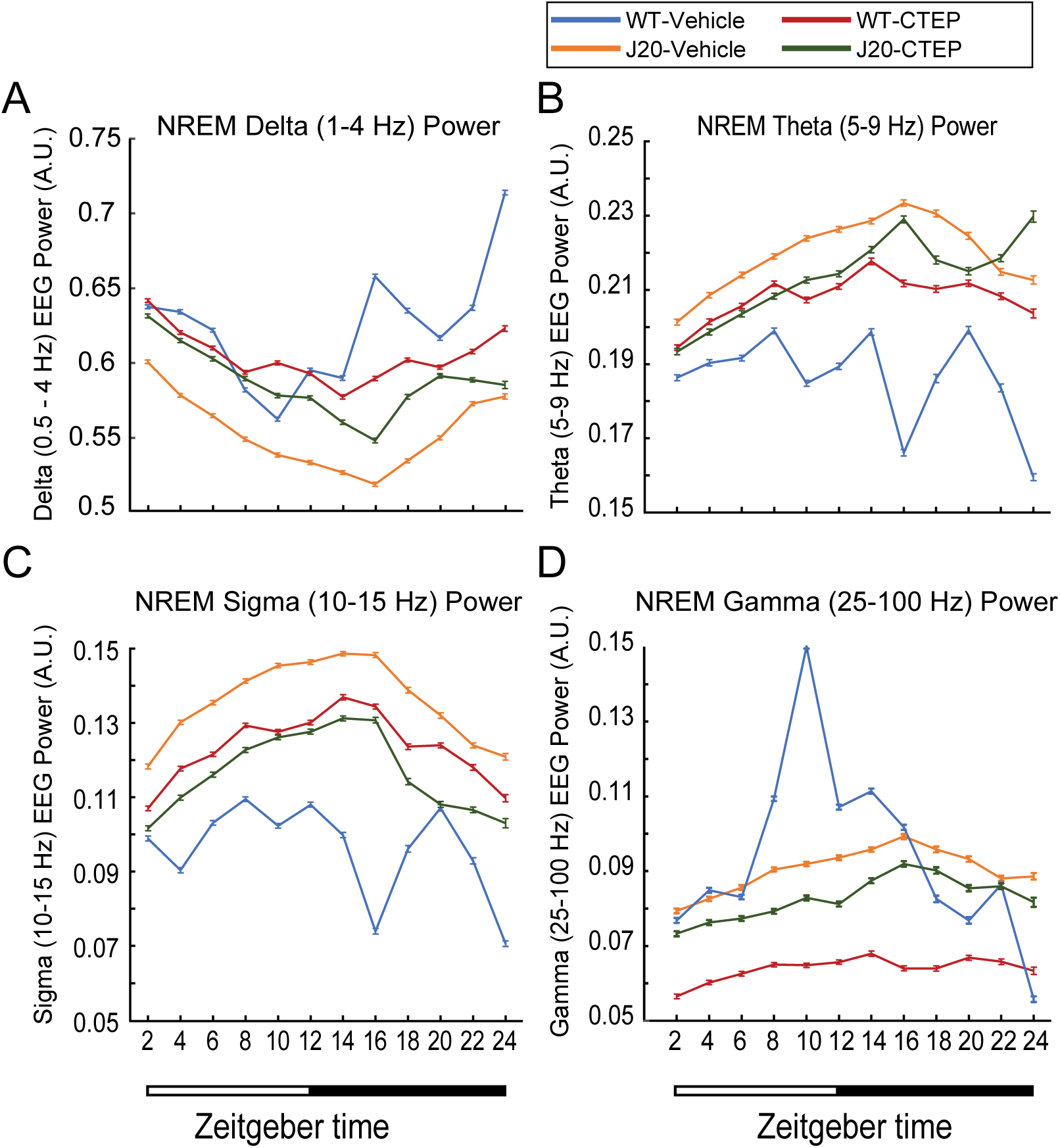
Power spectra of NREM sleep. Electroencephalographic power spectra of 4-second NREM sleep epochs were determined with fast-Fourier transform. Resulting power of delta (1-4 Hz), theta (5-9 Hz), sigma (10-15 Hz), and gamma (25-100 Hz) frequencies were isolated and normalized to the summed power of all frequency bands. Each 24-hour recording was divided into 2-hour segments, and manually scored NREM epochs within each bin were grouped. Here, normalized (A) delta, (B) theta, (C) sigma and (D) gamma powers are presented as mean ± 95% confidence interval (CI) for each treatment group: wild type (WT) treated with vehicle (n=3 mice for 3 days); J20 treated with vehicle (n=4 mice for 3 days); WT treated with CTEP (2-chloro-4-((2,5-dimethyl-1-(4-(trifluoromethoxy)phenyl)-1H-imidazol-4-yl)ethynyl)pyridine) (n=4 mice for 3 days), and J20 treated with CTEP (n=3 mice for 3 days). Non-overlapping CI bars indicate a significant difference (*P*<0.05). Lighting conditions are shown below the graph.

### Chronic mGluR_5_ Inhibition does not Significantly Reduce Aβ Levels

There was decreased (32%) plasma Aβ_1-40_ in J20 mice in response to fenobam that was not statistically significant by 2-way ANOVA [WT/vehicle 64.61±6.95, WT/fenobam 78.81±13.07, J20/vehicle 209.71±27.18, J20/fenobam 142.35±14.77; 2-way ANOVA: interaction *p*=0.031, F(1,23)=5.27; drug treatment *p*=0.15, F(1,23)=2.24; genotype, *p*<0.0001, F(1,23)=34.45]), and no other differences in Aβ_1-40_ or Aβ_1-42_ levels observed in plasma or brain for either strain treated with fenobam or CTEP (**Figure S7**). Chronic dosing with fenobam or CTEP did not affect mouse performance in passive avoidance or rotarod testing (**Figures S8 & S9**). Of note, we achieved low bioavailability of CTEP in the mice using established oral gavage dosing protocols. Based on published studies, we expected the dosing regimen to result in a minimal (trough level) drug exposure of 98±14 ng/mL in plasma and 215±28 ng/g in brain;^21, 27^ however, we achieved at least a 50-fold lower dose in both blood and brain (**Figure S10**). Poor bioavailability could be due a variety of factors (**Text S1)**.

## Discussion

AD is characterized by agitation and disruptions in activity and sleep typically observed in the evening, suggesting that diurnal dysfunction could serve as a disease biomarker.^28^ We employed actigraphy and EEG to determine rest-activity patterns and sleep phenotypes, respectively, under diurnal conditions in J20 AD mice and WT littermate controls to determine if rest-activity cycles are a reliable phenotype that can be pharmaceutically rescued in the mice as well as employed as a surrogate for EEG. Our major findings are that: (1) rest-activity rhythms are disrupted in J20 mice; and (2) sleep regulation is disrupted in J20 mice as evidenced by reduced NREM EEG delta power, which was partially rescued with CTEP. Low levels of CTEP bioavailability limit our ability to firmly conclude if CTEP has effects on sleep-activity rhythms. Since we did not record sleep EEG with fenobam, we cannot comment on its effects on sleep.

### J20 Mice

There are numerous mouse models available for the study of AD with many exhibiting altered sleep-wake states and diurnal rest-activity rhythms, albeit, there are variations in outcomes among the models.^29–34^ Our study is the first, to our knowledge, to directly compare diurnal rest-activity levels to sleep phenotypes in J20 mice. J20 mice are transgenic for the human amyloid protein precursor (hAPP) gene with the Swedish 670/671KM-NL and Indiana 717V-F double mutations under regulation by the PDGFβ chain promoter.^35^ The transgenic construct contains 70 bases of 5’-UTR, the cDNA, and the 3’-UTR up to the Sph1 site (base 3119 of APP_695_). The inclusion of flanking sequences in the transgenic construct is expected to affect posttranscriptional regulation of the *APP* gene and temporal and spatial expression of APP and metabolites. J20 mice are devoid of 3D6-immunoreative Aβ deposits at 2-4 months of age, but amyloid deposition can be observed in 50% of J20 mice by 5-7 months of age and in 100% of mice by 8-10 months (human equivalent, 42-50 years old).^35, 36^

### Actigraphy and EEG

Actigraphy provides an indirect measure of sleep-wake patterns and can determine gross shifts in rest-activity patterns. Actigraphy values are expected to be low during rest periods and high during active periods. J20 mice exhibit an altered rest-activity rhythm characterized by a 4-hour shift in the acrophase of peak activity, delays in activity onset and activity offset, and increased total activity during the dark cycle. Activity onset is a measure of the time of day in which the animals begin their most active period, and activity offset is a measure of when this active period ends. Since rodents are nocturnal, activity onset should begin at or around the start of the dark phase.^37^ Our findings are consistent with clinical studies showing later acrophase in AD patients.^38–40^ J20 mice are hyperactive in the open field and exhibit reduced anxiety.^41, 42^ Likewise, we found decreased habituation to the novel actigraphy environment.

EEG-based analysis showed minimal differences in time spent in NREM and REM sleep, but a profound decrease in NREM EEG-delta power in J20 mice. Specifically, with regard to REM, vehicle-treated J20 mice have a lower marginal mean (a weighted estimate of population means) of percent time spent in REM sleep from Zeitgeber time (ZT) 6-12 hours compared to vehicle-treated wild type animals, which is rescued by CTEP in J20 mice. According to the two-process model of sleep introduced by Borbély, sleep is regulated by both circadian and homeostatic mechanisms.^43^ The latter has been described as the pressure for sleep that grows during periods of wakefulness and is expunged by NREM sleep. Delta power is a commonly employed correlate of homeostatic sleep pressure, normally decreasing across the light period when mice are mainly resting and increasing with activity across the active dark period.^44, 45^ This relationship between activity and EEG delta power of NREM sleep is evidenced by substantial increases in delta (1-4 Hz) EEG power following brief (4 hours) total sleep deprivation in mice.^46^ Our findings agree with previous murine AD studies demonstrating a decrease in delta band EEG power, while activity of higher frequencies is increased.^47^ This apparent shift may be due to the large decreases in J20 NREM delta power, which has the largest influence on the power normalization, but it may also be indicative of hyperexcitability of neurons contributing to activity outside the typical on-off periods underlying high-amplitude, slow activity recorded at cortical surfaces during NREM sleep.^48^ Importantly, the increase in delta power occurs later in the subjective day, and it can be assumed that it increases in proportion to delayed waking duration and associated intensity of activity in J20 mice. It appears that CTEP treatment improves delta power (sleep pressure), although it reduces oscillatory amplitude in both WT and J20. Overall, the cyclic decay and accrual of delta power across the 24-hour period fits reasonably well with actigraphy, suggesting its viability as a substitute diagnostic tool for AD in place of invasive EEG-based methods. Taken together with delayed acrophase in locomotor activity observed by actigraphy during the dark phase, the phase-shifted NREM delta power may indicate perturbed function of the central pacemaker, affecting typical consolidation of sleep to subjectively appropriate times of the day.

### mGluR_5_ Inhibition

All of the currently approved drugs for the treatment of AD act on healthy neurons to compensate for lost acetylcholine activity in the case of cholinesterase inhibitors or to modulate NMDA receptor activity in the case of memantine. They improve cognitive ability for a year or less, but do not reduce Aβ accumulation or subsequent disease progression. The therapeutic potential of targeting mGluR_5_ in AD has been reviewed.^49^ *App* mRNA is a synaptic target for regulation by FMRP and mGluR_5_. Activation of mGluR_5_ signaling induces the release of the translational repressor FMRP from *App* mRNA and the subsequent synthesis of AβPP.^12^ Excessive AβPP production favors amyloidogenic processing and the production of Aβ. Aβ disrupts human NREM slow waves and related hippocampus-dependent memory consolidation.^50^ We have observed that treatment with the mGluR_5_ inhibitor CTEP partly rescued the sleep phenotype, which has not been previously reported to our knowledge. There is evidence that mGluR_5_ may have a modulatory role in the molecular machinery of sleep homeostasis.^51^ Thus, mGluR_5_ inhibitors may affect sleep-wake patterns but further study into the mechanism is required. Whether this translates to improved cognition remains to be determined.

Fenobam and CTEP are potent and highly selective noncompetitive inhibitors of mGluR_5._^27, 52, 53^ CTEP has a 30- to 100-fold higher *in vivo* potency compared to MPEP and fenobam and is 1,000-fold more selective for mGluR_5_ when compared to 103 molecular targets including all known mGluRs.^27^ Thus, if fenobam and/or CTEP are proven effective in reducing Aβ accumulation and the cognitive decline associated with AD, mGluR_5_ inhibitors could provide an alternative, orally administered treatment for AD, which lack the problems associated with antibody-based therapies. The dose of fenobam used herein (24 mg/kg/day) was calculated based on published rodent and human pharmacokinetic data. Phase I dose escalation trials showed safety and a lack of cognitive dysfunction in humans receiving up to 8-9 mg/kg/day fenobam for 3 weeks.^54^ Thus, the dose is 3-fold higher than that safely tested in humans, but far less than that safely tested in rats.^55^ Chronic dosing for 10-weeks at this dose resulted in no adverse side effects on weight gain or home cage behavior.^15^ As there are no reports of toxicity with the drug, we proposed to err on the side of over-dosing to ascertain fenobam effects on learning & memory and biomarker expression. CTEP is the first reported mGluR_5_ inhibitor with both a long half-life of approximately 18 hours and a high oral bioavailability, allowing chronic treatment with continuous receptor blockade with one dose every 48 hours in adult animals.^27^ Chronic treatment (2 mg/kg every 48 hours) inhibits mGluR_5_ with a receptor occupancy of 81% and rescues cognitive deficits in *Fmr1^KO^* mice.^21^ For this study, we dosed by oral gavage as published pharmacokinetic data by this method is available in other rodent models.^21, 27^

Treatment with mGluR_5_ inhibitors, fenobam or CTEP, did not rescue altered rest-activity profiles or affect mouse performance in rotarod or passive avoidance testing, although there were modest improvements in NREM delta power in CTEP-treated J20 mice. Oddly, oral gavage with vehicle shifts peak acrophase in WT mice. Specifically, in **Figure 3B** with oral gavage every 48 hours, average peak acrophase occurs at least 1 hour later in the WT mice, thus attenuating differences observed between WT and J20 in the absence of restraint/oral gavage (**Figure 1**). Statistically increased activity in J20 is observed at the end of the dark phase. This is a finding that we could not explain. The actigraphy and behavioral analyses involved chronic dosing with CTEP over 30 days whereas the EEG involved treatment for 1 week. Drug tolerance with mGluR_5_ inhibitors has been raised as an issue in failed FXS clinical trials.^56^ Consistent with chronic dosing studies of fenobam as a feed supplement in AD mice (Tg2576 and R1.40^HET^), and with chronic dosing of CTEP by oral gavage in FXS mice (*Fmr1^KO^*),^15, 21^ we observed normal weight gain, motor activity, grooming and home cage behavior with no adverse side effects. In contrast, genetic reduction of mGluR_5_ or chronic oral administration of CTEP rescues spatial learning deficits in APP_SWE_/PS1dE9 mice,^14, 17^ and BMS-984923 mGluR_5_ inhibitor treatment rescues memory deficits and synaptic depletion in APP_SWE_/PS1dE9 mice.^57^ We did not find genotype or drug dependent effects on learning & memory by passive avoidance in J20 mice.

### Study Limitations

Limitations of the study include poor bioavailability and possibly drug tolerance of the mGluR_5_ inhibitors, use of one AD mouse model, mice are nocturnal, and the oral gavage procedure shifts peak acrophase in WT mice. To begin to address these issues, future studies can include administration of drugs as feed supplements, testing the effects of drugs that modulate Aβ production such as β-secretase inhibitors, and determination of diurnal activity and sleep patterns in additional AD mouse models.

## Conclusion

In conclusion, sleep disturbances and behavioral symptoms are the main reasons to institutionalize patients with AD.^58, 59^ We observed disruptions in rest-activity rhythms and sleep in J20 mice, and sleep was partially rescued with one mGluR_5_ inhibitor. Chronic treatment with mGluR_5_ inhibitors did not rescue rest-activity rhythms in J20 mice, but altered dosing administration methods or alternative drugs may be effective and deserve further investigation. Actigraphy was a reasonable surrogate for EEG with the noted limitations. Overall, targeting sleep may be an avenue to delay the development and/or progression of AD.

## Acknowledgments

The authors thank Dr. Andrzej Dekundy (Merz Pharmaceuticals GmbH) and Dr. Lothar Lindemann (F. Hoffmann-La Roche) for advice on CTEP inhibitor solubility. This research was supported by NIA grant AG044714, the University of Wisconsin-Madison Alzheimer’s Disease Research Center (ADRC) grant P50-AG033514, the Clinical and Translational Science Award (CTSA) program through the National Center for Advancing Translational Sciences (NCATS) grant UL1TR002373, the Department of Defense (DOD) grant W81XWH-16-1-0082, and FRAXA Research Foundation.

## List of Abbreviations

AD: Alzheimer’s disease
ADRC: Alzheimer’s Disease Research Center
Aβ: amyloid-beta
AβPP: amyloid beta protein precursor
CTSA: Clinical and Translational Science Award
DOD: Department of Defense
DEA: diethylamine
EEG: electroencephalography
EMG: electromyography
FXS: fragile X syndrome
*hAPP*: human amyloid protein precursor
HPMC: hypromellose
mGluR_5_: metabotropic glutamate receptor 5
NCATS: National Center for Advancing Translational Sciences
NREM: non-rapid eye movement
REM: rapid eye movement
WT: wild type

## Ethics Statement

All mouse procedures were performed in accordance with NIH guidelines and an approved University of Wisconsin-Madison animal care protocol administered through their Institutional Animal Care and Use Committee (IACUC).

---

**Figure S1:**
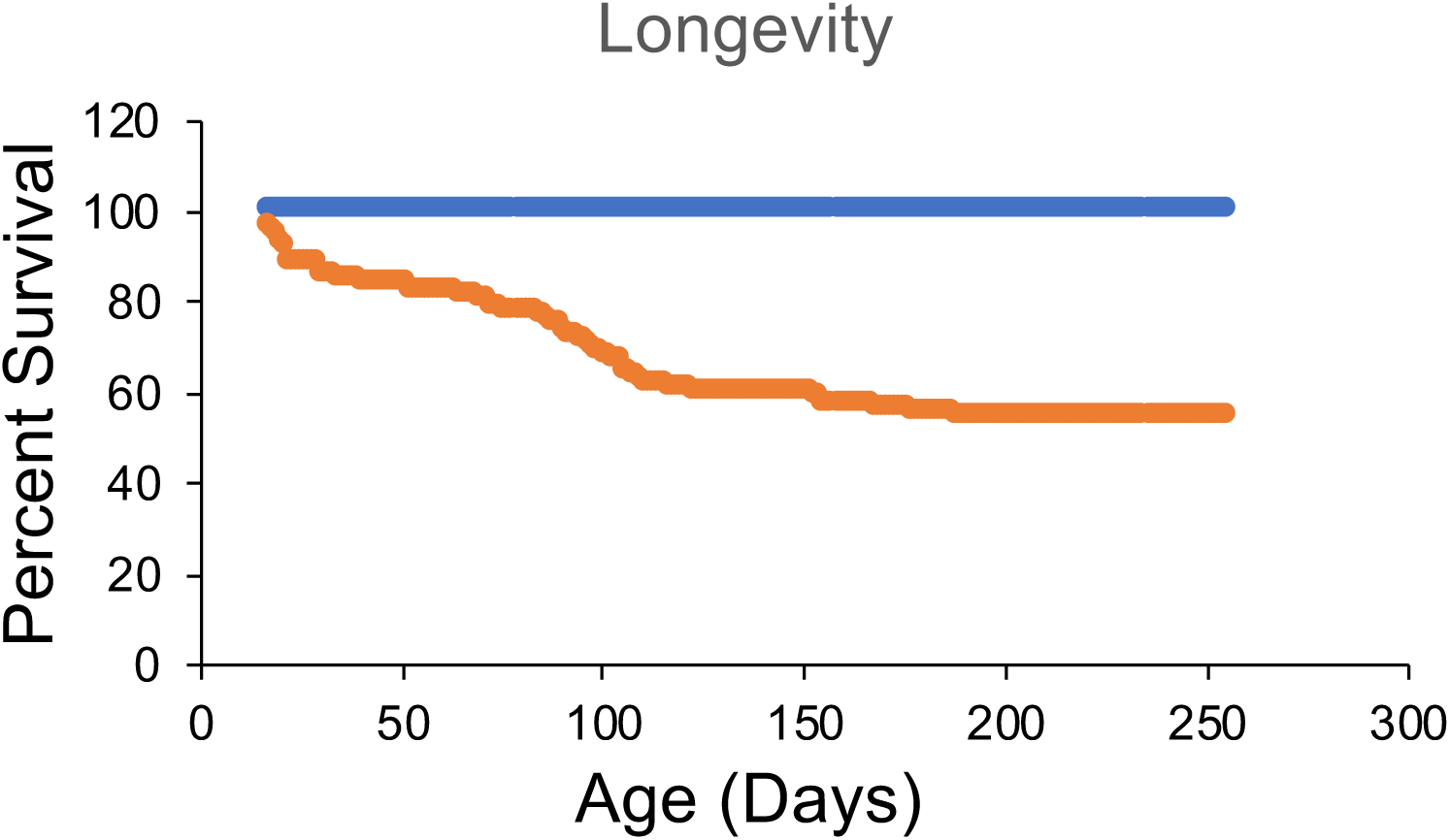
Mouse longevity of wild type (WT) and J20 littermates through 9 months of age. J20 mice (orange; n=114) exhibited a 40% premature death rate by 3 months of age compared to 0% premature deaths in WT mice (blue; n=123). Data for male mice is shown. Females exhibited a similar profile through 3 months of age (data not shown because females were not housed for the extended period of time shown for the males).

**Figure S2:**
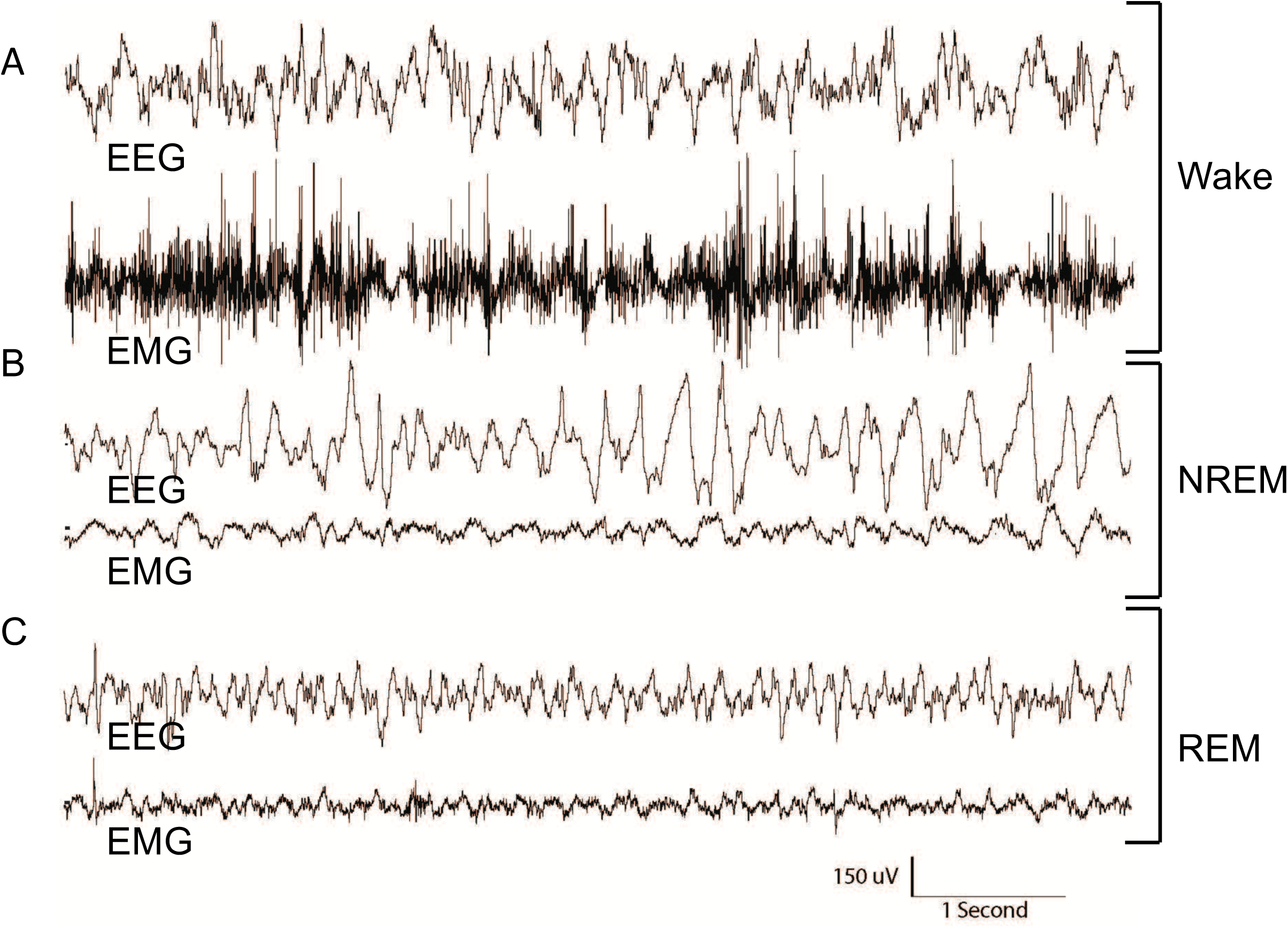
Representative EEG traces for sleep vigilance states. (A) Wakefulness is typically characterized by high-EMG amplitude. (B and C) Manual score of sleep necessitates a low EMG amplitude, where NREM is dominated by a slow, large amplitude delta power (1-4 Hz), while REM sleep contains lower amplitude waveforms with largely theta band (6-9 Hz) activity.

**Figure S3:**
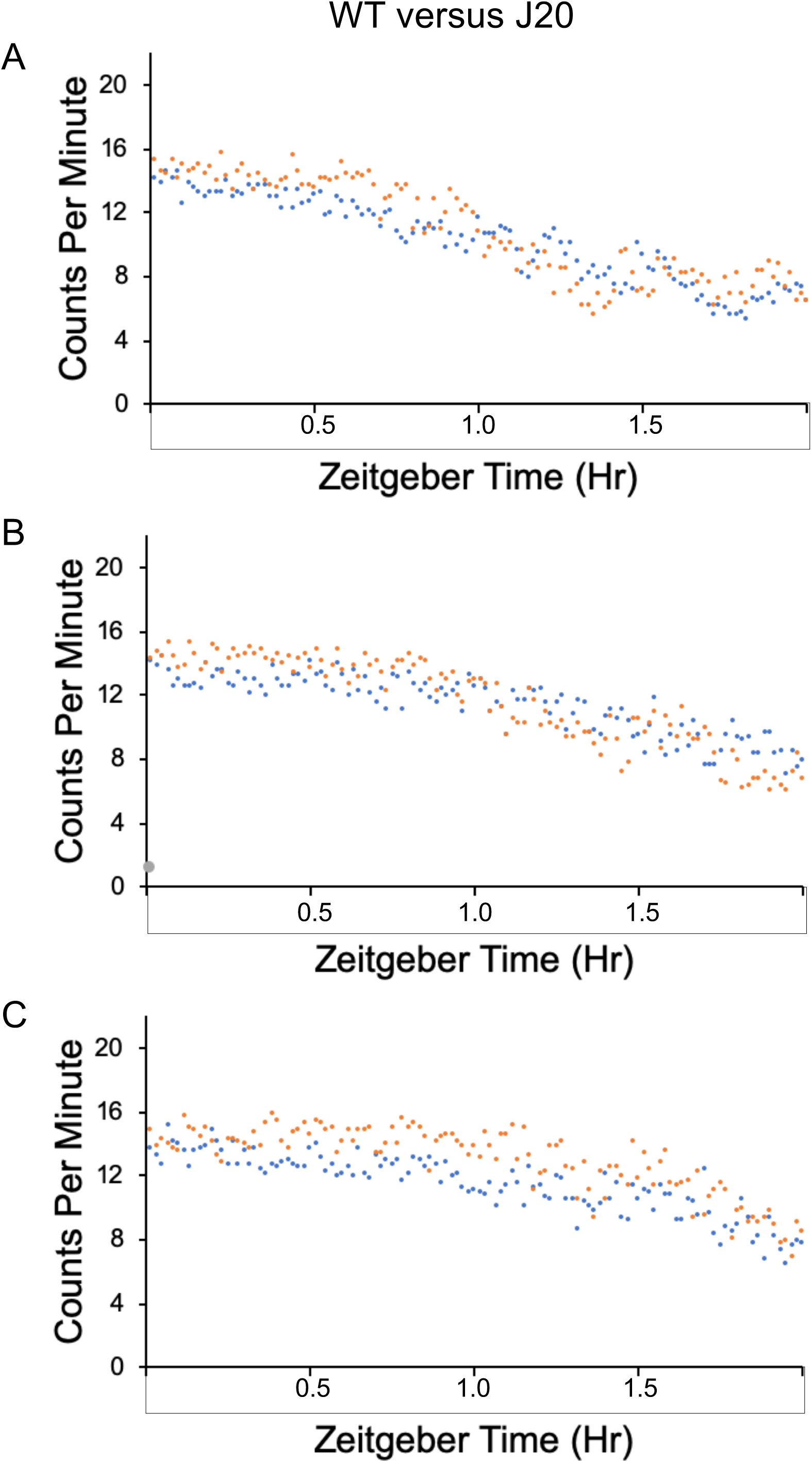
J20 mice exhibit decreased habituation. Habituation during the first 2 hours in the actigraphy chambers was assessed in 3 separate cohorts of wild type (WT) (blue) and J20 (orange) 8-month old mice (A=fall, B=winter, C=spring). Total activity counts (binned in 1 minute increments) were averaged for cohorts and plotted on the y-axis versus the first 120 minute time period on the x-axis. (A) Cohort consists of WT (n=15) and J20 (n=13). (B) Cohort consists of WT (n=12) and J20 (n=11). (C) Cohort consists of (n=12) and J20 (n=8).

**Figure S4:**
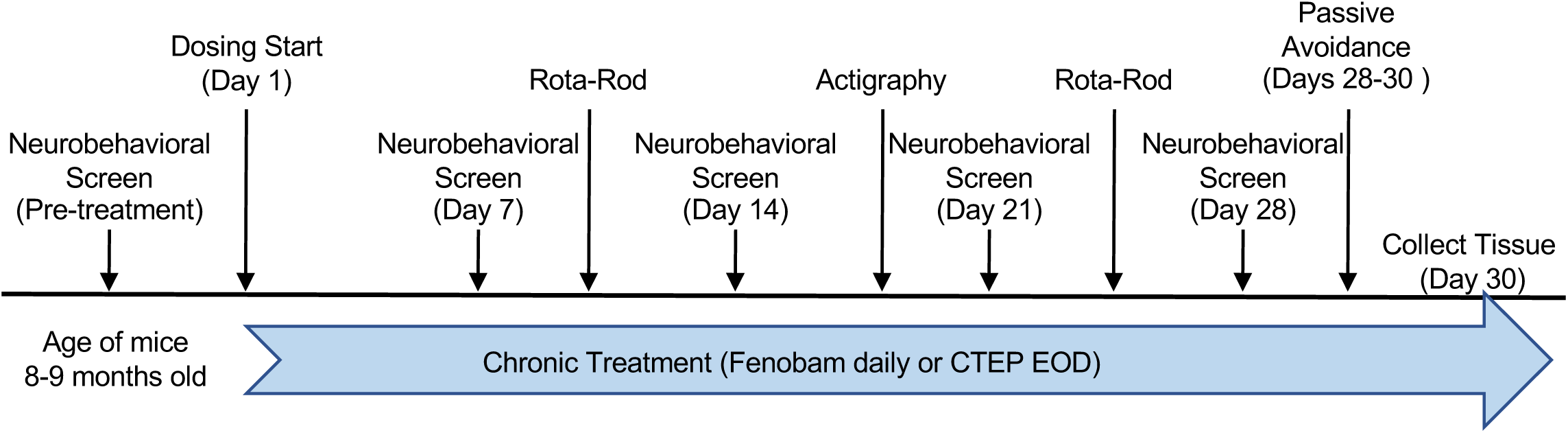
Dosing scheme and behavioral battery for wild type (WT) and J20 mice. The timeline for testing and dosing included a pre-treatment neurobehavioral screen at 8-9 months of age with follow-up neurobehavioral screens at 7, 14, 21 and 28 days dosing. Fenobam was dosed daily and CTEP every other day. Rotarod testing was conducted during weeks 2 & 4 of dosing, actigraphy during week 3 of dosing, and passive avoidance on dosing days 28-30. Tissue was collected on dosing day 30. All procedures were performed during the light cycle.

**Figure S5:**
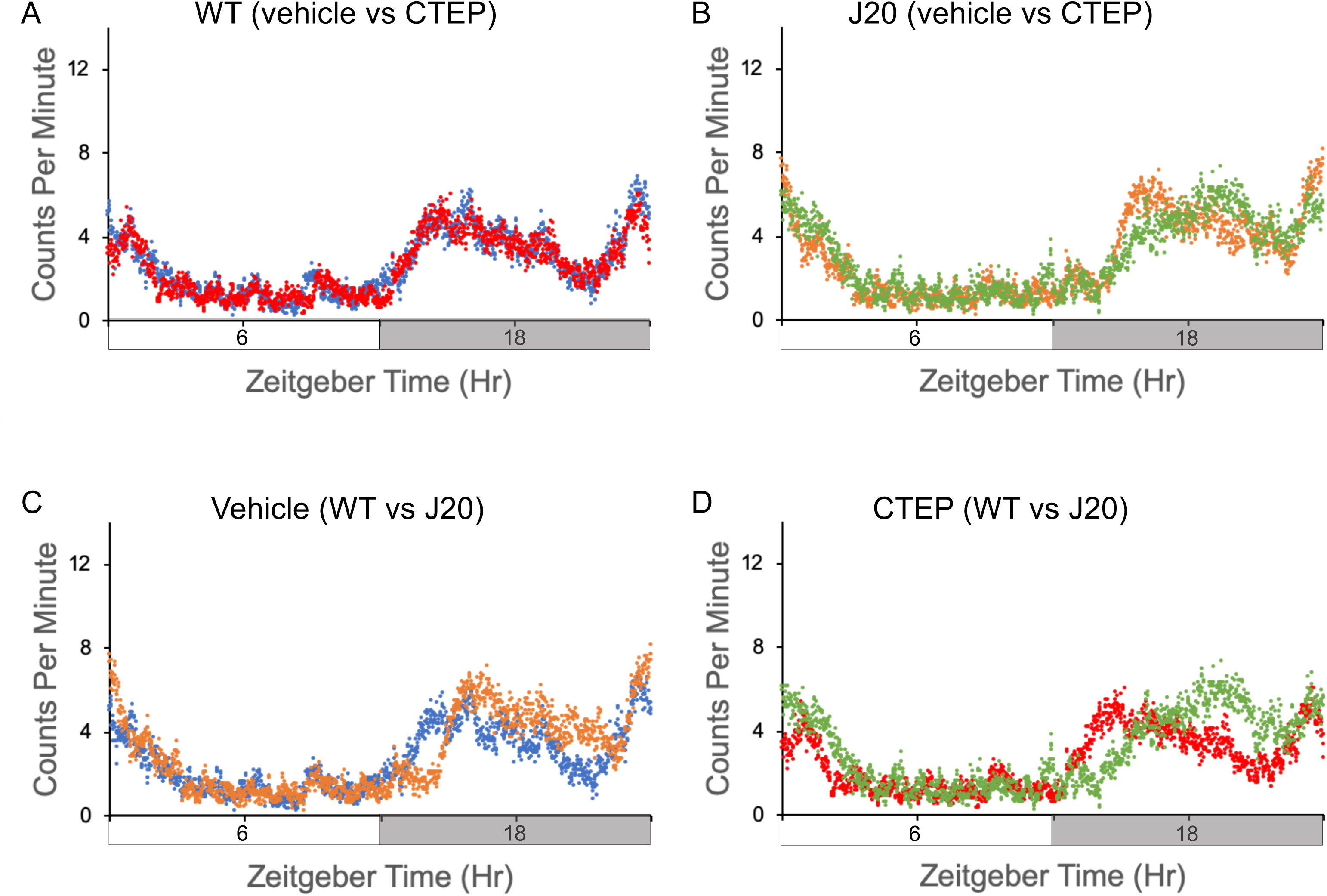
Diurnal activity levels in wild type (WT) and J20 mice in response to CTEP (2-chloro-4-((2,5-dimethyl-1-(4-(trifluoromethoxy)phenyl)-1H-imidazol-4-yl)ethynyl)pyridine). Activity counts were assessed in 16-19-month old WT and J20 mice after chronic treatment with vehicle or CTEP. Total activity counts (binned in 1 minute increments) were averaged over 7 days of readings for cohorts and plotted on the y-axis versus a 24 hour time period (in minutes). Time zero is “Lights On”. Cohorts consist of WT mice treated with vehicle (n=7, blue), J20 treated with vehicle (n=7, orange), WT treated with CTEP (n=9, red), and J20 treated with CTEP (n=6, green). (A) WT mice treated with vehicle (blue) versus CTEP (red). (B) J20 mice treated with vehicle (orange) versus CTEP (green). (C) Vehicle-treated WT (blue) versus J20 (orange). (D) CTEP-treated WT (red) versus J20 (green). 2-way ANOVA results: interaction *p*>0.9999, F(4317, 37440)=0.11; treatment (genotype and drug) *p*<0.0001, F(3, 37440)=12.34; time, *p*<0.0001, F(1439, 37440)=1.36.

**Figure S6:**
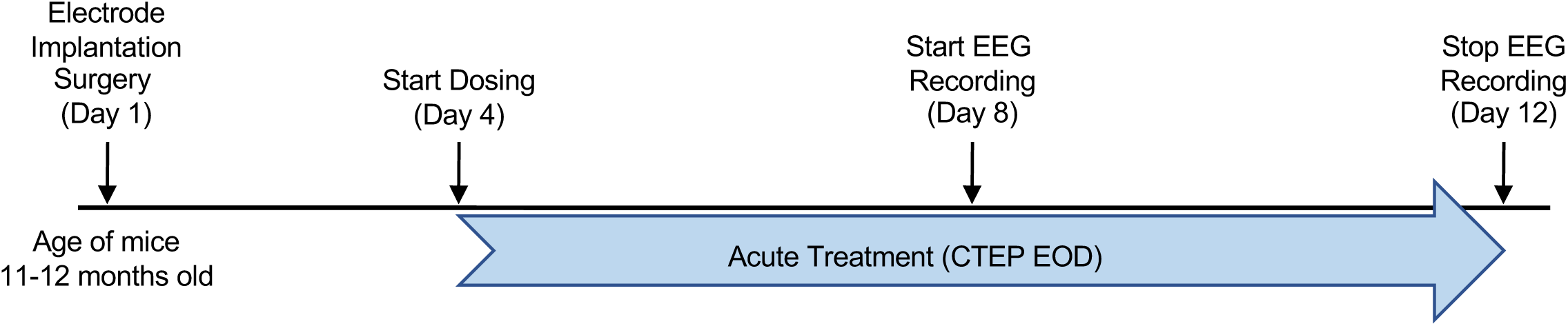
Dosing scheme for wild type (WT) and J20 mice for the electroencephalography (EEG) study. Mice (11-2 months old) were implanted with electrodes as described in the Methods on Day 1. After 3 days recovery from surgery (Day 4), mice received vehicle or CTEP treatment every other day (EOD). EEG recordings were collected from Day 8-12. Dosing continued EOD throughout the EEG recordings.

**Figure S7:**
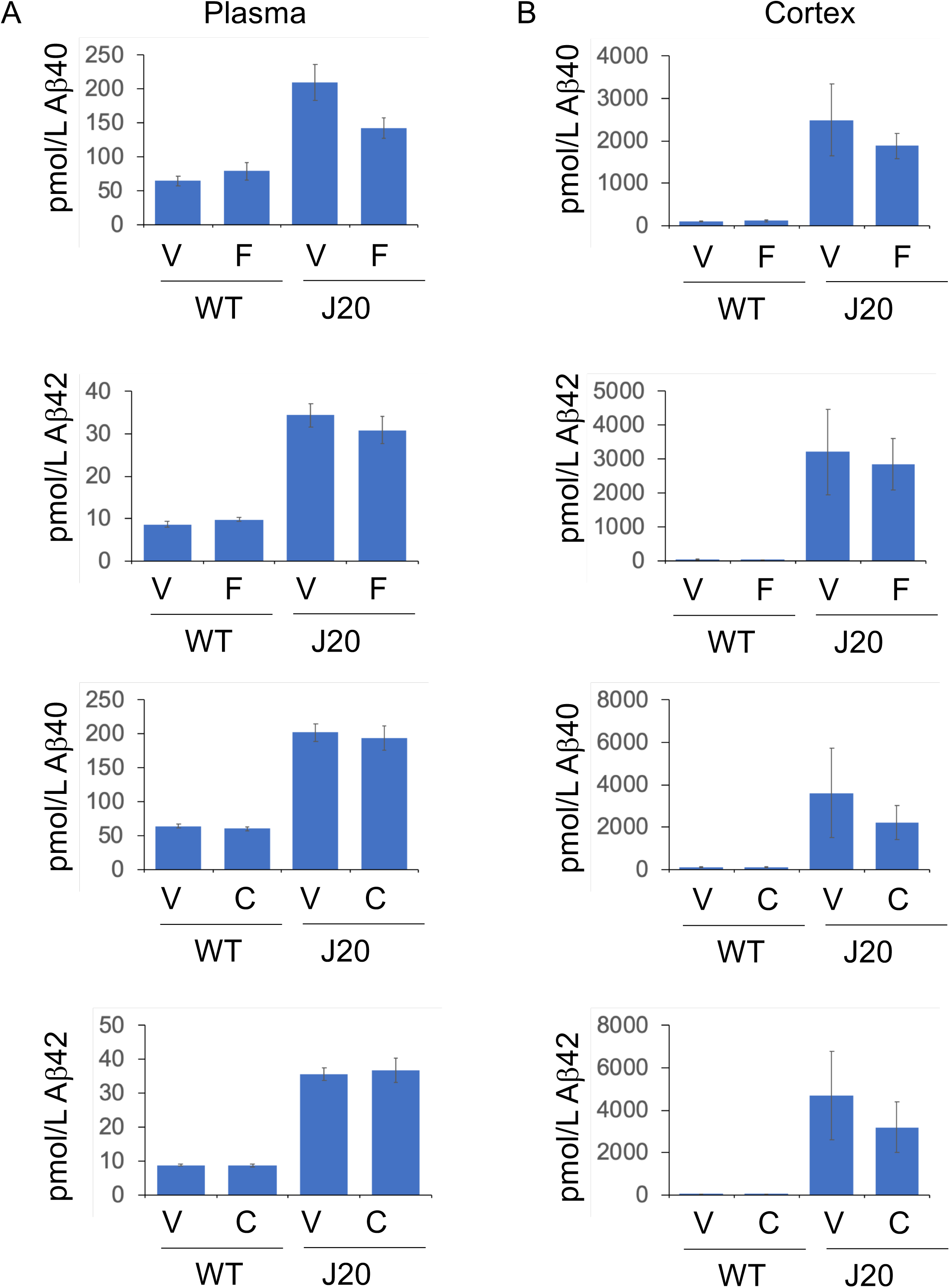
Amyloid-beta (Aβ) levels in response to metabotropic glutamate receptor 5 (mGluR_5_) inhibition. Aβ_1-40_ and Aβ_1-42_ levels were quantitated in (A) blood plasma and (B) brain (right cortex) from 8.5-10-month old WT and J20 mice by ELISA after chronic treatment with fenobam or CTEP (2-chloro-4-((2,5-dimethyl-1-(4-(trifluoromethoxy)phenyl)-1H-imidazol-4-yl)ethynyl)pyridine). Optical densities at 450 nm were converted to pmol/L, corrected for dilution factors, and plotted versus treatment/genotype conditions. Fenobam cohorts consisted of wild type (WT) mice treated with vehicle daily (V) (n=7), WT mice treated with fenobam daily (F) (n=7), J20 mice treated with vehicle daily (n=7), and J20 mice treated with fenobam daily (n=7). CTEP cohorts consisted of WT mice treated with vehicle every other day (EOD) (n=14), WT mice treated with CTEP EOD (n=12), J20 mice treated with vehicle EOD (n=9), and J20 mice treated with CTEP EOD (n=12). Error bars indicate standard error of the mean (SEM). 2-way ANOVA results: <u>plasma/Aβ40/fenobam</u>: interaction *p*=0.032, F(1,23)=5.2; drug treatment *p*=0.15, F(1,23)=2.2; genotype, *p*<0.0001, F(1,23)=34 <u>plasma/Aβ40/CTEP</u>: interaction *p*=0.86, F(1,43)=0.03; drug treatment *p*=0.58, F(1,43)=0.30; genotype, *p*<0.0001, F(1,43)=153 <u>cortex/Aβ40/fenobam</u>: interaction *p*=0.51, F(1,23)=0.45; drug treatment *p*=0.52, F(1,23)=0.42; genotype, *p*=0.0002, F(1,23)=19.83 <u>cortex/Aβ40/CTEP</u>: interaction *p*=0.44, F(1,43)=0.59; drug treatment *p*=0.45, F(1,43)=0.57; genotype, *p*=0.0037, F(1,43)=9.43 <u>plasma/Aβ42/fenobam</u>: interaction *p*=0.31, F(1,23)=1.08; drug treatment *p*=0.61, F(1,23)=0.27; genotype, *p*<0.0001, F(1,23)=108.6 <u>plasma/Aβ42/CTEP:</u> interaction *p*=0.77, F(1,43)=0.084; drug treatment *p*=0.79, F(1,43)=0.072; genotype, *p*<0.0001, F(1,43)=193.8 <u>cortex/Aβ42/fenobam:</u> interaction *p*=0.82, F(1,23)=0.052; drug treatment *p*=0.82, F(1,23)=0.055; genotype, *p*=0.0007, F(1,23)=15.24 <u>cortex/Aβ42/CTEP:</u> interaction *p*=0.47, F(1,43)=0.54; drug treatment *p*=0.47, F(1,43)=0.54; genotype, *p*=0.0004, F(1,43)=14.90

**Figure S8:**
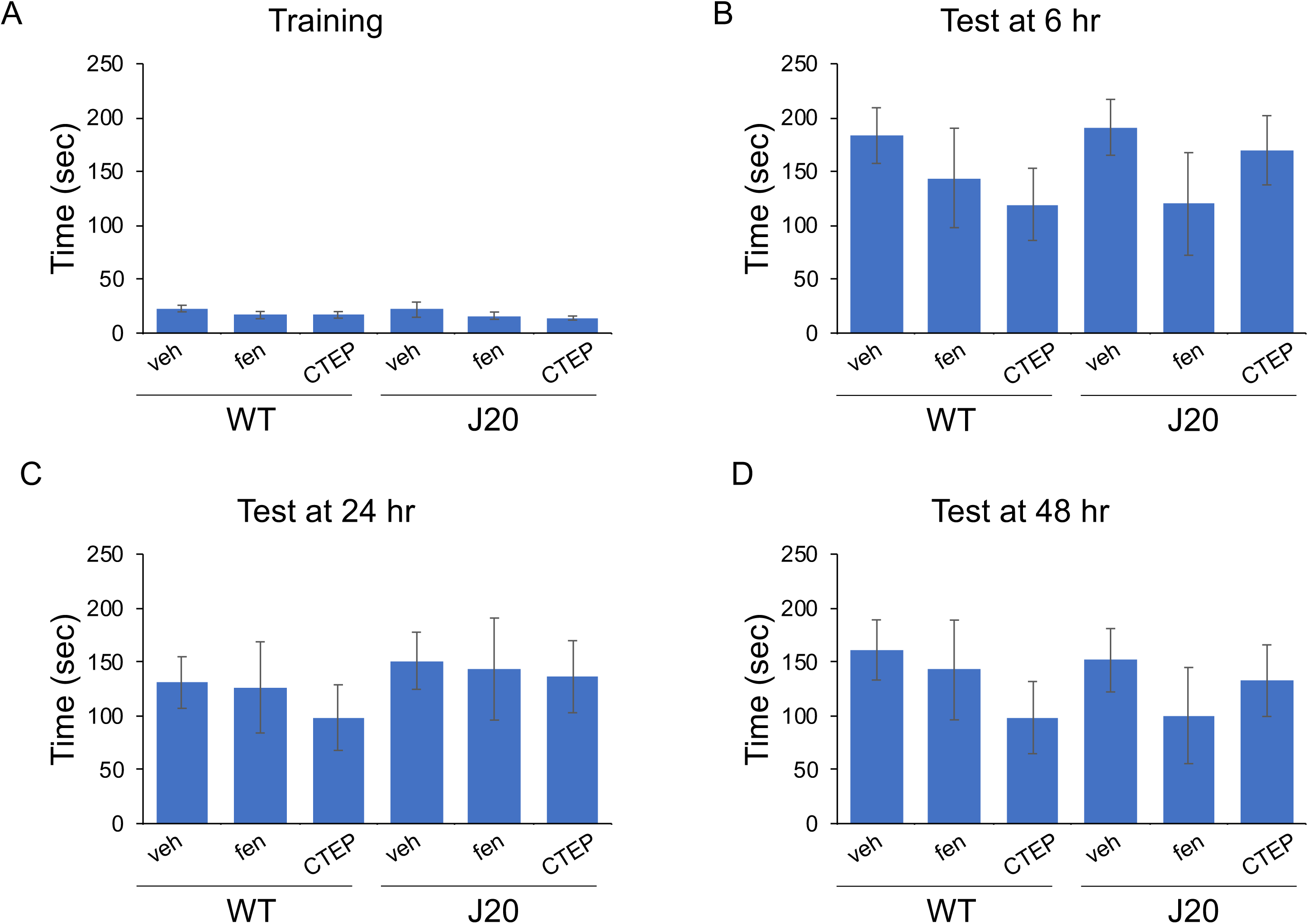
Learning and memory in response to metabotropic glutamate receptor 5 (mGluR_5_) inhibition. Learning and memory was assessed by passive avoidance testing. Mice were (A) trained with a 2-second 0.6 mA footshock on the dark side of a light-dark chamber and tested for learning and memory (B) 6 hours, (C) 24 hours, and (D) 48 hours later by latency time to enter the dark side. Neither genotype (WT = wild type; J20 = Alzheimer’s mice) nor treatment (veh = vehicle, fen = fenobam, CTEP = 2-chloro-4-((2,5-dimethyl-1-(4-(trifluoromethoxy)phenyl)-1H-imidazol-4-yl)ethynyl)pyridine) significantly affected learning & memory in the passive avoidance task.

**Figure S9:**
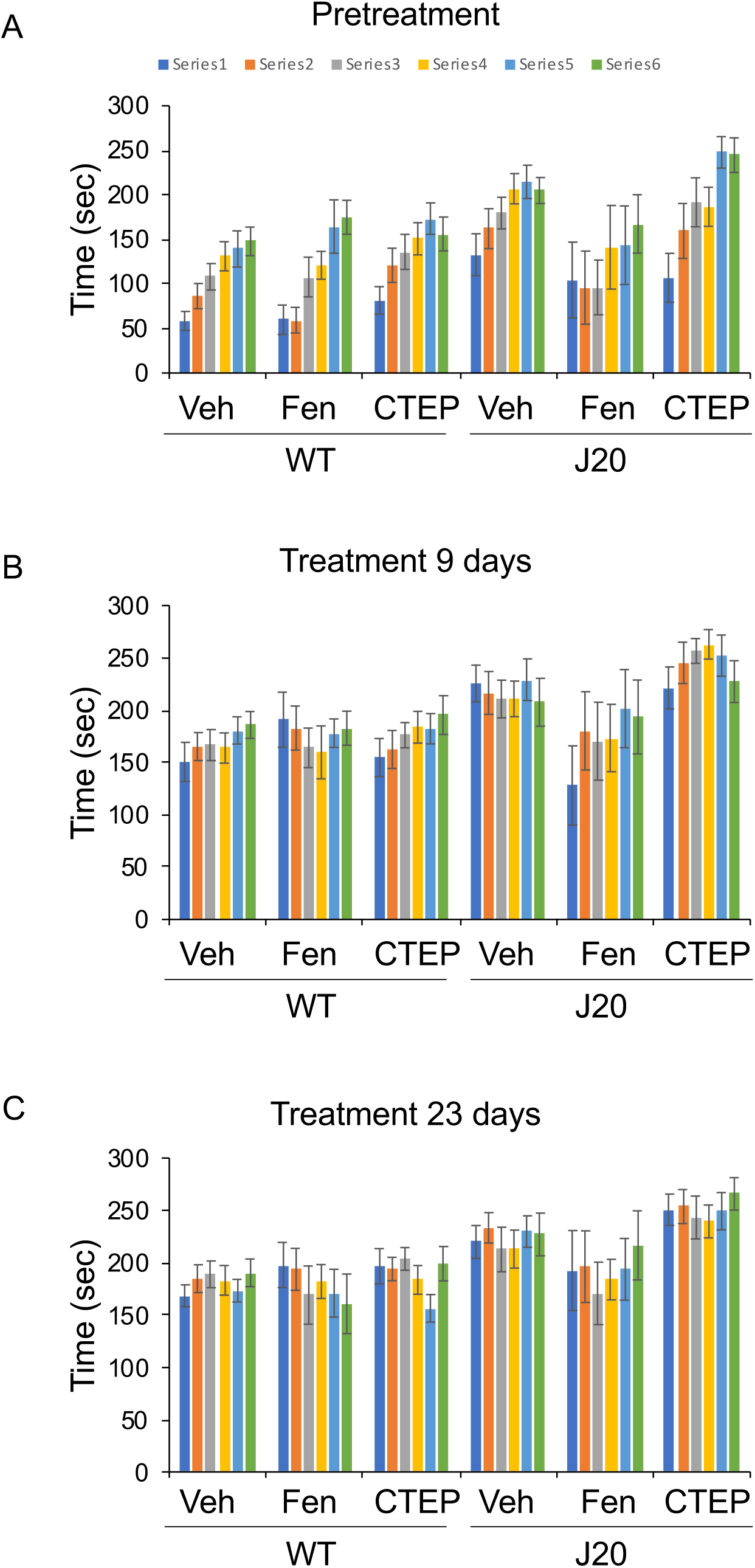
Motor coordination in response to metabotropic glutamate receptor 5 (mGluR_5_) inhibition. Motor coordination was assessed on the rotarod (A) pretreatment, (B) after 9 days treatment with veh (vehicle), Fen (fenobam) or CTEP (2-chloro-4-((2,5-dimethyl-1-(4-(trifluoromethoxy)phenyl)-1H-imidazol-4-yl)ethynyl)pyridine), and (C) after 23 days treatment with veh, Fen or CTEP. Six trials or series were run for each experiment with 4 trials on Day 1 and 2 trials on Day 2. Treatment did not impair motor coordination. All treatment groups exhibited increased motor learning with an increased number of trials during the pretreatment testing. All treatment groups maintained the maximal motor coordination ability attained during the pretreatment testing in subsequent testing after both 9 and 23 days of treatment.

**Figure S10:**
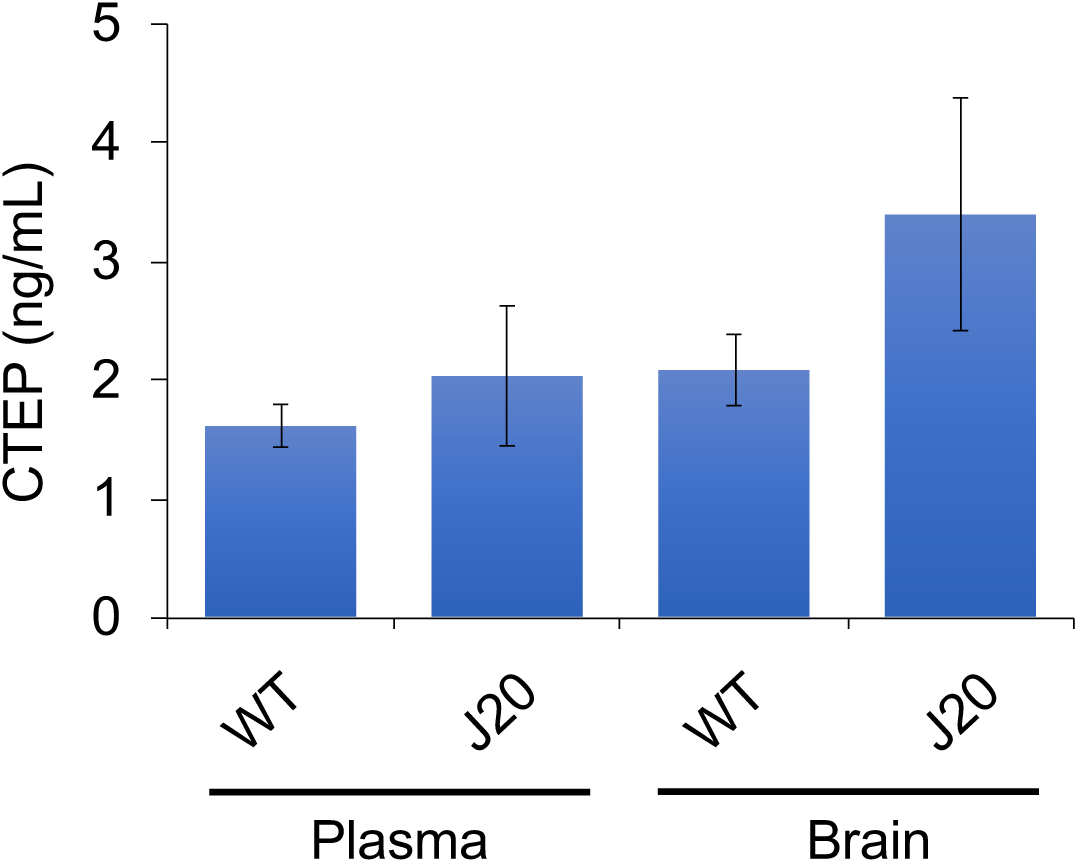
CTEP (2-chloro-4-((2,5-dimethyl-1-(4-(trifluoromethoxy)phenyl)-1H-imidazol-4-yl)ethynyl)pyridine) levels in blood plasma and brain after chronic dosing. CTEP levels were quantitated in blood plasma and brain cortical tissue from wild type (WT) (n=12) and J20 (n=11) mice (9-months old) after chronic drug dosing at 2 mg/kg by *Method 1*.

## Text S1

Upon consultation with pharmaceutical industry experts on dosing with mGluR_5_ inhibitors, we were advised that: (1) drug properties can differ dependent on the source of the compound, i.e. they prefer to use in-house prepared compounds versus commercially synthesized drugs; (2) differences in pharmacokinetics usually parallel differences in the physical property and/or solubility of the compound, (3) it may be advantageous to use a suspending agent such as HPMC; and (4) some of them have had great difficulty in reproducing pharmacokinetic results using published solvents and protocols. Thus, we modified the drug preparation protocol (*Method 2*) and tested the dose dependency (0.1, 0.5 and 2 mg/kg) of CTEP pharmacokinetics by oral gavage in WT mice (2-months old) with blood plasma collected 2.25 hours after dosing. Due to material transfer agreement issues, we continued to use a commercial source of CTEP. We found a clear dose response in drug levels with a single dose of CTEP by oral gavage [0.1 mg/kg: 4.2±2.2 ng/mL; 0.5 mg/kg: 30±1.3 ng/mL; 2 mg/kg: 128±19 ng/mL]. Thus, *Method 2* dosing was used for Figure 5 (aged mice). However, after 25 days chronic dosing with CTEP every other day (EOD) in aged WT and J20 cohorts, pharmacokinetic analysis of samples collected 1 hr after the last CTEP dose indicated 17.8±10.2 ng/mL CTEP in WT plasma (n=10), 14.3±21.9 ng/mL in WT brain (n=11), 12.9±8.8 ng/mL in J20 plasma (n=8), and 9.5±5.3 ng/mL in J20 brain (n=8). All were 6-23-fold lower than expected.

## Supplementary Tables

**Table S1:** Neurological Assessment Metrics (Fenobam) Mice underwent a pre-treatment evaluation of general fitness and grip strength and weekly assessments throughout dosing with fenobam. Fractions indicate the number of mice exhibiting a phenotype, for example, “No (7/7)” indicates that 7 out of a total of 7mice did not exhibit the listed phenotype and “Yes (1/7)” indicates that 1 mouse out of a total of 7 mice exhibited the listed phenotype.

| Table S1: Neurological Assessment Metrics (Fenobam) |  |  |  |  |  |  |
| --- | --- | --- | --- | --- | --- | --- |
|  |  |  | WT<br>Vehicle<br>n=7 | J20<br>Vehicle<br>n=7 | WT<br>Fenobam<br>n=8 | J20<br>Fenobam<br>n=6 |
| Body Weight (grams)<br>± SEM |  | start | 39.22±1.24 | 37.58±2.23 | 39.50±1.37 | 43.98±2.30 |
|  |  | 1 week | 37.00±1.05 | 34.93±2.12 | 38.00±1.39 | 42.11±2.36 |
|  |  | 2 weeks | 35.89±1.15 | 34.79±2.15 | 37.13±1.37 | 41.58±2.49 |
|  |  | 3 weeks | 34.76±1.34 | 34.80±1.77 | 36.16±1.32 | 41.94±2.67 |
|  |  | 4 weeks | 34.13±1.19 | 33.78±1.71 | 35.91±1.31 | 40.88±2.70 |
| General Observations |  |  | WT<br>Vehicle | J20<br>Vehicle | WT<br>Fenobam | J20<br>Fenobam |
| Tremor | Shaking<br>movements | start | No (7/7) | No (7/7) | No (8/8) | No (6/6) |
|  |  | 1 week | No (7/7) | No (7/7) | No (8/8) | No (6/6) |
|  |  | 2 weeks | No (7/7) | No (7/7) | No (8/8) | No (6/6) |
|  |  | 3 weeks | No (7/7) | No (7/7) | No (8/8) | No (6/6) |
|  |  | 4 weeks | No (7/7) | No (7/7) | No (8/8) | No (6/6) |
| Eyelid<br>Closure | Eyelids<br>closed<br>(cataracts) | start | No (7/7) | No (7/7) | No (6/7) | No (6/6) |
|  |  | 1 week | No (7/7) | No (7/7) | No (6/7) | No (6/6) |
|  |  | 2 weeks | No (6/7) | No (7/7) | No (6/7)<br>Yes (1/7) | No (6/6)<br>Yes (1/7) |
|  |  | 3 weeks | No (6/7) | No (7/7) | No (6/7)<br>Yes (1/7) | No (6/6)<br>Yes (1/7) |
|  |  | 4 weeks | No (6/7) | No (7/7) | No (6/7)<br>Yes (1/7) | No (6/6)<br>Yes (1/7) |
| Piloerection | Involuntary<br>erection or<br>bristling of<br>hairs | start | No (7/7) | No (7/7) | No (8/8) | No (6/6) |
|  |  | 1 week | No (7/7) | No (7/7) | No (8/8) | No (6/6) |
|  |  | 2 weeks | No (7/7) | No (7/7) | No (8/8) | No (5/6) |
|  |  | 3 weeks | No (7/7) | No (7/7) | No (8/8) | No (6/6) |
|  |  | 4 weeks | No (6/7) | No (7/7) | No (8/8) | No (6/6) |
| Lacrimation | Secretion of<br>tears | start | No (7/7) | No (7/7) | No (8/8) | No (6/6) |
|  |  | 1 week | No (7/7) | No (7/7) | No (8/8) | No (6/6) |
|  |  | 2 weeks | No (7/7) | No (7/7) | No (8/8) | No (6/6) |
|  |  | 3 weeks | No (7/7) | No (7/7) | No (8/8) | No (6/6) |
|  |  | 4 weeks | No (7/7) | No (7/7) | No (8/8) | No (6/6) |
| Salivation | Secreting<br>saliva | start | No (7/7) | No (7/7) | No (8/8) | No (6/6) |
|  |  | 1 week | No (7/7) | No (7/7) | No (8/8) | No (6/6) |
|  |  | 2 weeks | No (7/7) | No (7/7) | No (8/8) | No (6/6) |
|  |  | 3 weeks | No (7/7) | No (7/7) | No (8/8) | No (6/6) |
|  |  | 4 weeks | No (7/7) | No (7/7) | No (8/8) | No (6/6) |
| Dirty Coat | Coat is dirty | start | No (7/7) | No (7/7) | No (8/8) | No (6/6) |
|  |  | 1 week | No (7/7) | No (7/7) | No (8/8) | No (6/6) |
|  |  | 2 weeks | No (7/7) | No (7/7) | No (8/8) | No (6/6) |
|  |  | 3 weeks | No (7/7) | No (7/7) | No (8/8) | No (6/6) |
|  |  | 4 weeks | No (7/7) | No (7/7) | No (8/8) | No (6/6) |
| Grip<br>Strength | Mouse<br>allowed to<br>grip cage feed | start | 2.1±0.3 | 1.6±0.2 | 1.5±0.2 | 1.3±0.2 |
|  |  | 1 week | 1.1±0.1 | 1.4±0.2 | 1.4±0.2 | 1.2±0.2 |
|  |  | 2 weeks | 1.6±0.2 | 1.1±0.1 | 1.5±0.2 | 1.7±0.2 |
|  | bin wire while investigator gently pulls backwards horizontally:<br>0 = no grip,<br>1 = slight grip,<br>2 = moderate grip,<br>3 = active grip,<br>4 = unusually effective grip | **3 weeks** | 1.7±0.2 | 1.4±0.2 | 1.6±0.3 | 1.3±0.2 |
|  |  | **4 weeks** | 1.6±0.3 | 1.3±0.2 | 1.4±0.2 | 1.5±0.2 |
| **Reflexes** |  |  | **WT Vehicle** | **J20 Vehicle** | **WT Fenobam** | **J20 Fenobam** |
| Righting Reflex | Hold mouse by tail and turn onto back. Failure to right their body into an upright position within 5 seconds is scored as No. | **start** | Yes (7/7) | Yes (7/7) | Yes (8/8) | Yes (6/6) |
|  |  | **1 week** | Yes (7/7) | Yes (7/7) | Yes (8/8) | Yes (6/6) |
|  |  | **2 weeks** | Yes (7/7) | Yes (7/7) | Yes (8/8) | Yes (6/6) |
|  |  | **3 weeks** | Yes (7/7) | Yes (7/7) | Yes (8/8) | Yes (6/6) |
|  |  | **4 weeks** | Yes (7/7) | Yes (7/7) | Yes (8/8) | Yes (6/6) |
| Pinna Reflex | While the mouse is gently restrained, touch each ear canal lightly with the tip of a metal spatula. Monitor for ear retraction or head movement. Yes = movement in both ears. | **start** | Yes (6/7) | Yes (6/7) | Yes (7/8) | Yes (5/6) |
|  |  | **1 week** | Yes (7/7) | Yes (4/7) | Yes (6/8) | Yes (5/6) |
|  |  | **2 weeks** | Yes (7/7) | Yes (7/7) | Yes (8/8) | Yes (5/6) |
|  |  | **3 weeks** | Yes (7/7) | Yes (7/7) | Yes (8/8) | Yes (6/6) |
|  |  | **4 weeks** | Yes (7/7) | Yes (7/7) | Yes (8/8) | Yes (5/6) |
| Eyeblink Response | When a cotton swab is moved toward an open eye, the mouse blinks. Yes = at least one eye blinks. | **start** | Yes (6/7) | Yes (4/7) | Yes (7/8) | Yes (5/6) |
|  |  | **1 week** | Yes (5/7) | Yes (2/7) | Yes (4/8) | Yes (1/6) |
|  |  | **2 weeks** | Yes (7/7) | Yes (7/7) | Yes (7/8) | Yes (6/6) |
|  |  | **3 weeks** | Yes (7/7) | Yes (4/7) | Yes (5/8) | Yes (1/6) |
|  |  | **4 weeks** | Yes (7/7) | Yes (4/7) | Yes (7/8) | Yes (6/6) |

| Observations in Arena |  |  | WT Vehicle | J20 Vehicle | WT Fenobam | J20 Fenobam |
| --- | --- | --- | --- | --- | --- | --- |
| Body Posture | In home cage:<br>0 = no movement, totally flattened posture;<br>1 = signs of some ataxia, slightly flattened posture;<br>2 = elevated posture | **start** | 2 (7/7) | 2 (7/7) | 2 (/8/8) | 2 (/6/6) |
|  |  | **1 week** | 2 (7/7) | 2 (7/7) | 2 (/8/8) | 2 (/6/6) |
|  |  | **2 weeks** | 2 (7/7) | 2 (7/7) | 2 (/8/8) | 2 (/6/6) |
|  |  | **3 weeks** | 2 (7/7) | 2 (7/7) | 2 (/8/8) | 2 (/6/6) |
|  |  | **4 weeks** | 2 (7/7) | 2 (7/7) | 2 (/8/8) | 2 (/6/6) |
| Transfer Arousal | After transfer to arena:<br>0 = coma,<br>1 = slight,<br>2 = freeze & move,<br>3 = normal (momentary stop/freeze allowed),<br>4 = swift,<br>5 = maniac | **start** | 3 (7/7) | 3 (7/7) | 3 (8/8) | 3 (6/6) |
|  |  | **1 week** | 3 (7/7) | 3 (7/7) | 3 (8/8) | 3 (6/6) |
|  |  | **2 weeks** | 3 (7/7) | 3 (7/7) | 3 (8/8) | 3 (6/6) |
|  |  | **3 weeks** | 3 (7/7) | 3 (7/7) | 3 (8/8) | 3 (6/6) |
|  |  | **4 weeks** | 3 (7/7) | 3 (7/7) | 3 (8/8) | 3 (6/6) |
| Tail Elevation | After transfer to arena: 0 = dragging,<br>1 = rattling,<br>2 = normal | **start** | 2 (7/7) | 2 (7/7) | 2 (/8/8) | 2 (/6/6) |
|  |  | **1 week** | 2 (7/7) | 2 (7/7) | 2 (/8/8) | 2 (/6/6) |
|  |  | **2 weeks** | 2 (7/7) | 2 (7/7) | 2 (/8/8) | 2 (/6/6) |
|  |  | **3 weeks** | 2 (7/7) | 2 (7/7) | 2 (/8/8) | 2 (/6/6) |
|  |  | **4 weeks** | 2 (7/7) | 2 (7/7) | 2 (/8/8) | 2 (/6/6) |

**Table S2:** Neurological Assessment Metrics (CTEP) Mice underwent a pre-treatment evaluation of general fitness and grip strength and weekly assessments throughout dosing with CTEP.

| Table S2: Neurological Assessment Metrics (CTEP) |  |  |  |  |  |  |
| --- | --- | --- | --- | --- | --- | --- |
|  |  |  | WT<br>Vehicle<br>n=14 | J20<br>Vehicle<br>n=9 | WT<br>CTEP<br>n=12 | J20<br>CTEP<br>n=12 |
| Body Weight (grams)<br>± SEM |  | start | 37.18±1.50 | 33.42±2.44 | 36.59±0.67 | 32.13±1.64 |
|  |  | 1 week | 36.67±1.42 | 32.57±2.41 | 35.90±0.64 | 30.94±1.51 |
|  |  | 2 weeks | 36.53±1.42 | 32.52±2.26 | 35.47±0.55 | 31.41±1.47 |
|  |  | 3 weeks | 36.66±1.69 | 32.37±2.14 | 35.48±0.74 | 31.73±1.41 |
|  |  | 4 weeks | 36.12±1.59 | 32.25±2.13 | 34.99±0.59 | 31.88±1.52 |
| General Observations |  |  | WT<br>Vehicle | J20<br>Vehicle | WT<br>CTEP | J20<br>CTEP |
| Tremors | Shaking movements | start | No (14/14) | No (9/9) | No (12/12) | No (12/12) |
|  |  | 1 week | No (14/14) | No (9/9) | No (12/12) | No (12/12) |
|  |  | 2 weeks | No (14/14) | No (9/9) | No (12/12) | No (12/12) |
|  |  | 3 weeks | No (14/14) | No (9/9) | No (12/12) | No (12/12) |
|  |  | 4 weeks | No (14/14) | No (9/9) | No (12/12) | No (12/12) |
| Eyelid<br>Closure | Eyelids closed<br>(cataracts) | start | No (14/14)<br>Yes (1/14) | No (9/9)<br>Yes (1/9) | No (11/12) | No (12/12) |
|  |  | 1 week | No (14/14)<br>Yes (1/14) | No (9/9)<br>Yes (1/9) | No (11/12) | No (12/12) |
|  |  | 2 weeks | No (14/14)<br>Yes (1/14) | No (9/9)<br>Yes (1/9) | No (11/12) | No (12/12) |
|  |  | 3 weeks | No (14/14)<br>Yes (1/14) | No (9/9)<br>Yes (1/9) | No (11/12) | No (12/12) |
|  |  | 4 weeks | No (14/14)<br>Yes (1/14) | No (9/9)<br>Yes (1/9) | No (11/12) | No (12/12) |
| Piloerection | Involuntary erection or<br>bristling of hairs | start | No (14/14) | No (9/9) | No (11/12) | No (12/12) |
|  |  | 1 week | No (14/14) | No (9/9) | No (11/12) | No (12/12) |
|  |  | 2 weeks | No (14/14) | No (9/9) | No (12/12) | No (12/12) |
|  |  | 3 weeks | No (14/14) | No (8/9) | No (12/12) | No (12/12) |
|  |  | 4 weeks | No (14/14) | No (9/9) | No (12/12) | No (12/12) |
| Lacrimation | Secretion of tears | start | No (14/14) | No (9/9) | No (12/12) | No (12/12) |
|  |  | 1 week | No (14/14) | No (9/9) | No (12/12) | No (12/12) |
|  |  | 2 weeks | No (14/14) | No (9/9) | No (12/12) | No (12/12) |
|  |  | 3 weeks | No (14/14) | No (9/9) | No (12/12) | No (12/12) |
|  |  | 4 weeks | No (14/14) | No (9/9) | No (12/12) | No (12/12) |
| Salivation | Secreting saliva | start | No (14/14) | No (9/9) | No (12/12) | No (12/12) |
|  |  | 1 week | No (14/14) | No (9/9) | No (12/12) | No (12/12) |
|  |  | 2 weeks | No (14/14) | No (9/9) | No (12/12) | No (12/12) |
|  |  | 3 weeks | No (14/14) | No (9/9) | No (12/12) | No (12/12) |
|  |  | 4 weeks | No (14/14) | No (9/9) | No (12/12) | No (12/12) |
| Dirty Coat | Coat is dirty | start | No (14/14) | No (9/9) | No (11/12) | No (12/12) |
|  |  | 1 week | No (14/14) | No (9/9) | No (12/12) | No (12/12) |
|  |  | 2 weeks | No (14/14) | No (9/9) | No (12/12) | No (12/12) |
|  |  | 3 weeks | No (14/14) | No (9/9) | No (12/12) | No (12/12) |
|  |  | 4 weeks | No (14/14) | No (9/9) | No (12/12) | No (12/12) |
| Grip<br>Strength | Mouse allowed to grip<br>cage feed bin wire while | start | 1.9±0.2 | 1.4±0.2 | 1.6±0.2 | 1.5±0.2 |
|  |  | 1 week | 1.4±0.1 | 1.4±0.2 | 1.5±0.2 | 1.6±0.3 |
|  |  | 2 weeks | 1.3±0.1 | 1.1±0.1 | 1.3±0.2 | 1.3±0.2 |
|  | investigator gently pulls backwards horizontally:<br>0= no grip, 1= slight grip<br>2 = moderate grip, 3 = active grip, 4 = unusually effective grip | **3 weeks** | 1.3±0.1 | 1.0±0.0 | 1.3±0.1 | 1.4±0.2 |
|  |  | **4 weeks** | 1.2±0.1 | 1.0±0.0 | 1.3±0.1 | 1.1±0.1 |
| **Reflexes** |  |  | **WT Vehicle** | **J20 Vehicle** | **WT CTEP** | **J20 CTEP** |
| Righting Reflex | Hold mouse by tail and turn onto back. Failure to right their body into an upright position within 5 seconds is scored as No. | **start** | Yes (14/14) | Yes (9/9) | Yes (12/12) | Yes (12/12) |
|  |  | **1 week** | Yes (14/14) | Yes (9/9) | Yes (12/12) | Yes (12/12) |
|  |  | **2 weeks** | Yes (14/14) | Yes (9/9) | Yes (12/12) | Yes (12/12) |
|  |  | **3 weeks** | Yes (14/14) | Yes (9/9) | Yes (12/12) | Yes (12/12) |
|  |  | **4 weeks** | Yes (14/14) | Yes (9/9) | Yes (12/12) | Yes (12/12) |
| Pinna Reflex | While the mouse is gently restrained, touch each ear canal lightly with the tip of a metal spatula. Monitor for ear retraction or head movement. Yes = movement in both ears. | **start** | Yes (13/14) | Yes (8/9) | Yes (10/12) | Yes (11/12) |
|  |  | **1 week** | Yes (13/14) | Yes (6/8) | Yes (11/12) | Yes (11/11) |
|  |  | **2 weeks** | Yes (14/14) | Yes (9/9) | Yes (11/12) | Yes (12/12) |
|  |  | **3 weeks** | Yes (14/14) | Yes (9/9) | Yes (12/12) | Yes (12/12) |
|  |  | **4 weeks** | Yes (12/14) | Yes (7/9) | Yes (10/12) | Yes (10/12) |
| Eyeblink Response | When a cotton swab is moved toward an open eye, the mouse blinks. Yes = at least one eye blinks. | **start** | Yes (12/14) | Yes (7/9) | Yes (10/12) | Yes (9/12) |
|  |  | **1 week** | Yes (11/14) | Yes (7/9) | Yes (10/12) | Yes (5/11) |
|  |  | **2 weeks** | Yes (9/14) | Yes (5/9) | Yes (10/12) | Yes (6/12) |
|  |  | **3 weeks** | Yes (13/14) | Yes (8/9) | Yes (12/12) | Yes (11/12) |
|  |  | **4 weeks** | Yes (14/14) | Yes (8/9) | Yes (12/12) | Yes (10/12) |
| **Observations in Arena** |  |  | **WT Vehicle** | **J20 Vehicle** | **WT CTEP** | **J20 CTEP** |
| Body Posture | In home cage: 0 = no movement, totally flattened posture; 1 = signs of some ataxia, slightly flattened posture; 2 = elevated posture | **start** | 2 (14/14) | 2 (8/9)<br>1 (1/9) | 2 (12/12) | 2 (12/12) |
|  |  | **1 week** | 2 (14/14) | 2 (9/9) | 2 (12/12) | 2 (12/12) |
|  |  | **2 weeks** | 2 (14/14) | 2 (9/9) | 2 (12/12) | 2 (12/12) |
|  |  | **3 weeks** | 2 (14/14) | 2 (9/9) | 2 (12/12) | 2 (12/12) |
|  |  | **4 weeks** | 2 (14/14) | 2 (9/9) | 2 (12/12) | 2 (12/12) |
| Transfer Arousal | After transfer to arena: 0 = coma, 1 = slight, 2 = freeze & move, 3 = normal (momentary stop/freeze allowed), 4 = swift, 5 = maniac | **start** | 3 (14/14) | 3 (9/9) | 3 (12/12) | 3 (12/12) |
|  |  | **1 week** | 3 (14/14) | 3 (9/9) | 3 (12/12) | 3 (12/12) |
|  |  | **2 weeks** | 3 (14/14) | 3 (9/9) | 3 (12/12) | 3 (12/12) |
|  |  | **3 weeks** | 3 (14/14) | 3 (9/9) | 3 (12/12) | 3 (12/12) |
|  |  | **4 weeks** | 3 (14/14) | 3 (9/9) | 3 (12/12) | 3 (12/12) |
| Tail Elevation | After transfer to arena: 0 = dragging,1 = rattling, 2 = normal | **start** | 2 (14/14) | 2 (7/9)<br>0 (2/9) | 2 (12/12) | 2 (12/12) |
|  |  | **1 week** | 2 (14/14) | 2 (9/9) | 2 (12/12) | 2 (12/12) |
|  |  | **2 weeks** | 2 (14/14) | 2 (9/9) | 2 (12/12) | 2 (12/12) |
|  |  | **3 weeks** | 2 (14/14) | 2 (9/9) | 2 (12/12) | 2 (12/12) |
|  |  | **4 weeks** | 2 (14/14) | 2 (9/9) | 2 (12/12) | 2 (12/12) |

**Table S3:** Mean Normalized NREM EEG Power Values by Group and Time. Marginal means±95% confidence interval from a mixed-effect ANOVA evaluating NREM EEG power for delta (0.5-4 Hz), theta (5-9 Hz), sigma (10-15 Hz), and gamma (25-100 Hz) frequency bands in J20 an WT mice with and without CTEP administration across 12, 2-hour bins beginning at lights-on.

| Table S3: Mean Normalized NREM EEG Power Values by Group and Time |  |  |  |  |  |  |  |  |
| --- | --- | --- | --- | --- | --- | --- | --- | --- |
| Delta (0.5-4 Hz) |  |  |  |  | Theta (5-9 Hz) |  |  |  |
| WT |  | J20 |  |  | WT |  | J20 |  |
| Vehicle | CTEP | Vehicle | CTEP | Time bin | Vehicle | CTEP | Vehicle | CTEP |
| 0.638±0.0013 | 0.642±0.0013 | 0.601±0.0013 | 0.632±0.0014 | 1 | 0.186±0.0008 | 0.194±0.0007 | 0.201±0.0008 | 0.193±0.0008 |
| 0.634±0.0014 | 0.620±0.0012 | 0.578±0.0012 | 0.615±0.0014 | 2 | 0.190±0.0008 | 0.201±0.0008 | 0.209±0.0006 | 0.199±0.0008 |
| 0.622±0.0012 | 0.610±0.0012 | 0.565±0.0012 | 0.603±0.0014 | 3 | 0.192±0.0007 | 0.206±0.0007 | 0.214±0.0007 | 0.204±0.0008 |
| 0.582±0.0013 | 0.594±0.0012 | 0.549±0.0012 | 0.590±0.0013 | 4 | 0.199±0.0007 | 0.212±0.0007 | 0.219±0.0007 | 0.208±0.0008 |
| 0.562±0.0013 | 0.600±0.0012 | 0.539±0.0012 | 0.578±0.0013 | 5 | 0.185±0.0007 | 0.207±0.0007 | 0.224±0.0007 | 0.213±0.0008 |
| 0.595±0.0013 | 0.593±0.0012 | 0.533±0.0013 | 0.577±0.0013 | 6 | 0.189±0.0008 | 0.211±0.0007 | 0.226±0.0008 | 0.214±0.0008 |
| 0.590±0.0014 | 0.577±0.0015 | 0.527±0.0012 | 0.560±0.0014 | 7 | 0.199±0.0008 | 0.218±0.0008 | 0.229±0.0007 | 0.221±0.0008 |
| 0.658±0.0015 | 0.590±0.0014 | 0.519±0.0014 | 0.548±0.0015 | 8 | 0.166±0.0009 | 0.212±0.0009 | 0.233±0.0007 | 0.229±0.0009 |
| 0.635±0.0018 | 0.602±0.0015 | 0.535±0.0016 | 0.577±0.0018 | 9 | 0.186±0.001 | 0.210±0.0009 | 0.231±0.0009 | 0.218±0.0011 |
| 0.617±0.0017 | 0.597±0.0014 | 0.550±0.0015 | 0.591±0.0016 | 10 | 0.199±0.001 | 0.212±0.0008 | 0.225±0.0009 | 0.215±0.0009 |
| 0.637±0.0017 | 0.608±0.0015 | 0.573±0.0014 | 0.589±0.0015 | 11 | 0.184±0.001 | 0.208±0.0009 | 0.215±0.0008 | 0.219±0.0009 |
| 0.714±0.0016 | 0.623±0.0020 | 0.578±0.0018 | 0.585±0.0025 | 12 | 0.160±0.0009 | 0.204±0.001 | 0.213±0.0011 | 0.230±0.0015 |

| Sigma (10-15 Hz) |  |  |  | Time bin | Gamma (25-100 Hz) |  |  |  |
| --- | --- | --- | --- | --- | --- | --- | --- | --- |
| WT |  | J20 |  |  | WT |  | J20 |  |
| Vehicle | CTEP | Vehicle | CTEP |  | Vehicle | CTEP | Vehicle | CTEP |
| 0.099±0.0006 | 0.107±0.0006 | 0.119±0.0006 | 0.102±0.0007 | 1 | 0.077±0.0006 | 0.056±0.0006 | 0.079±0.0006 | 0.073±0.0006 |
| 0.091±0.0007 | 0.118±0.0006 | 0.130±0.0005 | 0.110±0.0006 | 2 | 0.085±0.0007 | 0.060±0.0006 | 0.083±0.0005 | 0.076±0.0006 |
| 0.103±0.0006 | 0.122±0.0005 | 0.136±0.0005 | 0.116±0.0006 | 3 | 0.083±0.0006 | 0.063±0.0005 | 0.086±0.0005 | 0.077±0.0006 |
| 0.110±0.0006 | 0.130±0.0005 | 0.141±0.0005 | 0.123±0.0006 | 4 | 0.109±0.0006 | 0.065±0.0005 | 0.090±0.0005 | 0.079±0.0006 |
| 0.103±0.0006 | 0.128±0.0006 | 0.146±0.0005 | 0.126±0.0006 | 5 | 0.150±0.0006 | 0.065±0.0005 | 0.092±0.0005 | 0.083±0.0006 |
| 0.108±0.0007 | 0.130±0.0006 | 0.147±0.0006 | 0.128±0.0006 | 6 | 0.107±0.0007 | 0.066±0.0006 | 0.094±0.0006 | 0.081±0.0006 |
| 0.100±0.0007 | 0.137±0.0007 | 0.149±0.0005 | 0.131±0.0006 | 7 | 0.111±0.0007 | 0.068±0.0007 | 0.096±0.0005 | 0.087±0.0007 |
| 0.074±0.0007 | 0.135±0.0006 | 0.148±0.0006 | 0.131±0.0007 | 8 | 0.102±0.0007 | 0.064±0.0006 | 0.099±0.0006 | 0.092±0.0007 |
| 0.096±0.0009 | 0.124±0.0007 | 0.139±0.0007 | 0.114±0.0009 | 9 | 0.083±0.0009 | 0.064±0.0007 | 0.096±0.0007 | 0.090±0.0006 |
| 0.107±0.0008 | 0.124±0.0006 | 0.132±0.0007 | 0.108±0.0008 | 10 | 0.077±0.0008 | 0.067±0.0006 | 0.093±0.0007 | 0.085±0.0007 |
| 0.093±0.0008 | 0.118±0.0007 | 0.124±0.0007 | 0.107±0.0007 | 11 | 0.086±0.0008 | 0.066±0.0007 | 0.088±0.0006 | 0.086±0.0007 |
| 0.071±0.0007 | 0.110±0.0009 | 0.121±0.0008 | 0.103±0.0012 | 12 | 0.056±0.0007 | 0.063±0.0009 | 0.089±0.0009 | 0.082±0.0012 |

